# Phytochemical Modulation of Astrocyte A1/A2 Polarization and Hepcidin-Associated Iron Dysregulation in LPS-Driven Neuroinflammation

**DOI:** 10.64898/2026.05.14.725062

**Authors:** Masaki Kaneko, Chi-Fang Hsu, Cheng-Ta Tsai, Shion Osana, Takako Fujii, Susumu Ito, Katsuhiko Hata

## Abstract

**Background/Objectives:** Neuroinflammation-driven iron dysregulation and neurotoxic astrocyte polarization are increasingly recognized as interconnected pathological mechanisms in neurodegenerative diseases. Systemic inflammation triggered by strenuous exercise or infection can engage the central nervous system and astrocytic inflammatory responses and perturb iron homeostasis; however, targeted nutritional strategies to counteract these processes remain limited. Inflamate® is a multi-component botanical supplement comprising boswellic acids, astilbin, xanthohumol, and cinnamaldehyde, each with documented anti-inflammatory properties. However, whether this combined formulation can modulate the inflammatory–iron metabolic axis and astrocyte phenotypic polarization remains unexplored. This study aimed to investigate the effects of Inflamate® on LPS-induced pro-inflammatory gene expression, iron metabolism-related gene regulation, and A1/A2 astrocyte phenotypic polarization in mouse astrocytes.

**Methods:** Mouse astrocytes (AWT) were pre-treated with Inflamate® (0.0375 μg/mL) or DMSO vehicle for 24 h, followed by lipopolysaccharide (LPS; 1 μg/mL) stimulation for an additional 24 h. The non-cytotoxic working concentration was determined by morphological assessment, CCK-8 cell viability, and LDH cytotoxicity assays. Expression of 14 target genes spanning pro-inflammatory mediators (*NOS2, IL6, C3, COX2, PLA2g15, SOCS3*), iron metabolism regulators (*FTH1, Hepcidin, TFRC, SLC40A1, RGMa, RGMb*), and astrocyte polarization markers (*S100A10, GFAP*) was quantified by qRT-PCR.

**Results:** Under normal culture conditions, Inflamate® did not significantly alter the expression of any target gene except S100A10, confirming the absence of baseline cytotoxicity or transcriptional homeostatic perturbation. Upon LPS stimulation, Inflamate® selectively suppressed NOS2 (approximately 64% reduction, p < 0.0001), IL6 (approximately 37% reduction, p < 0.0001), and C3 (approximately 47% reduction, p < 0.0001), while COX2, PLA2g15, and SOCS3 remained unaffected. Concurrently, Inflamate® significantly reduced LPS-induced Hepcidin expression to approximately 17% of the control level (p < 0.05) and attenuated FTH1 upregulation (p < 0.01), without altering the expression of iron transporters (TFRC, SLC40A1) or BMP-SMAD pathway components (RGMa, RGMb). Furthermore, Inflamate® upregulated the neuroprotective A2 marker S100A10 under both basal (p < 0.05) and LPS-stimulated conditions (p < 0.01), while the general reactivity marker GFAP remained unchanged.

**Conclusions:** Inflamate® exerts a selective, multi-target modulatory effect at the transcriptional level in LPS-stimulated astrocytes, encompassing suppression of the iNOS–NO and IL-6 signaling axes, attenuation of inflammation-driven hepcidin–ferritin iron dysregulation via the IL-6–STAT3 pathway, and promotion of a phenotypic shift from neurotoxic A1 toward neuroprotective A2 astrocyte polarization. Given that the IL-6–JAK-STAT3–hepcidin axis is also activated during exercise-induced systemic inflammation, these findings suggest that Inflamate® may represent a targeted nutritional strategy for preserving CNS iron homeostasis and supporting neuroprotective astrocyte function in both neurodegenerative and exercise-related neuroinflammatory contexts. Further validation in in vivo neurodegenerative and exercise models, including protein-level analyses, is warranted to confirm these transcriptional findings.

## 1. Introduction

Inflammation-driven dysregulation of iron homeostasis is increasingly recognized as a critical interface between innate immunity and cellular metabolic balance[1, 2]. In the central nervous system (CNS), astrocytes are the most abundant glial cells and play essential roles in maintaining the brain microenvironment, including the regulation of iron metabolism, neurotransmitter recycling, and blood–brain barrier integrity[3–5]. Under pathological conditions, astrocytes undergo a process termed reactive astrogliosis, adopting distinct functional phenotypes that profoundly influence neuronal survival and disease progression[6]. Notably, systemic inflammation associated with strenuous or prolonged exercise has been suggested to increase blood–brain barrier permeability and may propagate peripheral inflammatory signals—including elevated IL-6 and TNF-α—into the CNS, potentially activating glial inflammatory responses that share mechanistic features with infection-driven neuroinflammation[7]. Moreover, prolonged strenuous exercise can promote gut-derived endotoxemia, with elevated circulating lipopolysaccharide levels reported in endurance athletes[8]. These observations provide a physiological motivation for investigating the IL-6–JAK-STAT3– hepcidin axis in the context of glial neuroinflammation and suggest that mechanistic insights derived from LPS-based cellular models may offer a relevant, if not directly equivalent, framework for understanding exercise-associated neuroinflammatory processes.

Upon exposure to lipopolysaccharide (LPS), a potent endotoxin derived from Gram-negative bacteria, the activation of nuclear factor kappa B (NF-κB) signaling in astrocytes triggers the robust production of pro-inflammatory mediators. Among these, inducible nitric oxide synthase (iNOS/NOS2) and interleukin-6 (IL-6) are two parallel downstream effectors of NF-κB that contribute to neuronal injury through distinct but complementary mechanisms [9–11]. NOS2-derived nitric oxide (NO) generates reactive nitrogen species that cause oxidative and nitrosative damage to neurons and oligodendrocytes[12, 13], while IL-6 activates the Janus kinase–signal transducer and activator of transcription 3 (JAK-STAT3) pathway, further amplifying the inflammatory cascade[14, 15]. Suppressor of cytokine signaling 3 (SOCS3) serves as a negative feedback regulator of this cascade, modulating the magnitude and duration of IL-6-driven signaling.

Critically, the IL-6–JAK-STAT3 axis directly upregulates HAMP transcription, the gene encoding hepcidin, the master regulator of iron homeostasis [16–18]. Hepcidin exerts its function primarily by binding to ferroportin (FPN1/SLC40A1), the sole known cellular iron exporter, thereby promoting its internalization and degradation [19]. This process leads to intracellular iron sequestration and a compensatory upregulation of ferritin heavy chain 1 (FTH1), the principal intracellular iron-storage protein [19–21]. In the brain, inflammation-induced elevation of hepcidin can cause a 40-fold increase in local hepcidin levels, and astrocyte- and microglia-derived hepcidin downregulates neuronal ferroportin, thereby promoting iron retention in the CNS microenvironment [22]. While FTH1 upregulation initially serves a cytoprotective function by sequestering labile iron and preventing Fenton chemistry-mediated generation of reactive oxygen species (ROS) [21, 22], sustained inflammation-driven iron accumulation may overwhelm this protective capacity [23, 24]. Under chronic inflammatory conditions, ferritinophagy—the NCOA4-mediated autophagic degradation of ferritin—can release stored iron back into the labile iron pool, paradoxically exacerbating oxidative damage and promoting ferroptosis[25–27]. This inflammation–iron retention axis is therefore considered a potential therapeutic target for mitigating secondary neuronal injury[28, 29]. In addition to the IL-6–STAT3 axis, hepcidin transcription is regulated by the bone morphogenetic protein–mothers against decapentaplegic homolog (BMP-SMAD) signaling pathway, in which repulsive guidance molecules A and B (RGMa and RGMb) function as essential BMP co-receptors[30, 31]. Although RGMa and RGMb have been extensively studied in the context of neuronal development and axon guidance, emerging evidence suggests that they may also participate in hepcidin regulation under inflammatory conditions [32–36].

Emerging evidence has further revealed that reactive astrocytes can be broadly classified into neurotoxic A1 and neuroprotective A2 phenotypes[6, 37–39]. Although it is increasingly recognized that astrocyte reactive states exist along a transcriptional continuum rather than as discrete binary categories [103, 104], the A1/A2 framework nonetheless provides a useful operational scaffold for interpreting the directional shift of key phenotypic markers—particularly C3 and S100A10—under defined inflammatory conditions, and is therefore retained as a conceptual framework in the present study. A1 astrocytes, induced by inflammatory signals such as LPS-activated microglial factors (TNF-α, IL-1α, and complement component C1q), are characterized by the upregulation of complement component C3 and contribute to neuronal and oligodendrocyte death[37, 40–42]. In contrast, A2 astrocytes, marked by the expression of the S100 calcium-binding protein A10 (S100A10), promote neuronal survival, synapse repair, and tissue remodeling [38, 43, 44]. The balance between A1 and A2 polarization states is a critical determinant of neurological outcomes following inflammatory insults, and pharmacological strategies that shift this balance toward the A2 phenotype hold considerable therapeutic promise [45, 46]. Notably, recent studies have demonstrated that anti-inflammatory interventions can simultaneously downregulate A1 markers (C3, iNOS) and upregulate A2 markers (S100A10, Arg1), indicating that these phenotypic markers are reciprocally regulated [47–50].In exercise physiology, it has been reported that repeated high-intensity exercise induces a persistent inflammatory environment in the central nervous system that promotes the polarization of neurotoxic A1-type astrocytes; this response may contribute to exercise-induced central fatigue and impaired neurological recovery[51]. In contrast, physical exercise itself shifts astrocyte polarization toward the neuroprotective A2 phenotype, with enhanced myelin debris clearance and improved cognitive function in a rat model of chronic cerebral hypoperfusion[52]. Therefore, strategies that promote the neuroprotective A2 phenotype may be relevant not only for neurodegenerative diseases but also for maintaining neurological health in physically active populations.

In the field of sports nutrition, dietary and botanical interventions[53] targeting exercise-induced inflammatory signaling have attracted considerable attention as potential modulators of neuroinflammation and iron dysregulation[54]. Inflamate® is a multi-component dietary supplement comprising four plant-derived bioactive ingredients: boswellic acids from Boswellia serrata resin, astilbin from Engelhardtia roxburghiana (Huangqi) leaves, xanthohumol from hop (Humulus lupulus) extract, and cinnamaldehyde from cinnamon (Cinnamomum cassia) bark[55, 56]. Individually, these compounds have demonstrated anti-inflammatory properties by suppressing NF-κB activation, iNOS expression, and pro-inflammatory cytokine release in various in vitro and in vivo models [57–62]. In particular, boswellic acids have been shown to inhibit NF-κB-mediated transcription of iNOS and TNF-α, and to exert neuroprotective effects by attenuating microglial and astrocyte activation[63, 64]. However, whether their combined formulation can modulate the inflammatory–iron metabolic axis and astrocyte polarization in the context of LPS-driven neuroinflammation remains largely unexplored.

The present study aimed to investigate the effects of Inflamate® on LPS-induced inflammatory gene expression, regulation of iron metabolism-related genes, and astrocyte phenotypic polarization in mouse astrocytes. We hypothesized that the anti-inflammatory properties of Inflamate® would attenuate LPS-driven NOS2 and IL-6 expression, thereby reducing inflammation-associated induction of hepcidin and ferritin, while concurrently promoting a shift from the neurotoxic A1 toward the neuroprotective A2 astrocyte phenotype. Elucidating these mechanisms at the cellular level may provide a mechanistic foundation for the rational application of phytochemical-based nutritional strategies in exercise-related neuroinflammatory contexts[65].

## 2. Materials and Methods

### 2.1. Cell Culture

Mouse astrocyte cell line (AWT) (RCB5681) were obtained from RIKEN BioResource Research Center (RIKEN BRC, Tsukuba, Japan) through the National Bio-Resource Project of the Ministry of Education, Culture, Sports, Science and Technology (MEXT), Japan. Cells were cultured under standard conditions (37 °C in a humidified atmosphere containing 5% CO₂) in high-glucose Dulbecco’s Modified Eagle Medium (DMEM; FUJIFILM Wako Pure Chemical Corporation, Osaka, Japan) supplemented with 10% heat-inactivated fetal bovine serum (FBS; Thermo Fisher Scientific, Waltham, MA, USA) and 100 μg/mL penicillin–streptomycin solution (FUJIFILM Wako Pure Chemical Corporation). Cell counts were determined using a Cell Counter model R1 (Olympus Corporation, Tokyo, Japan).

### 2.2. Preparation of Inflamate® and Experimental Design

#### 2.2.1. Preparation of Inflamate® Stock Solution

Inflamate® (boswellic acids from Boswellia serrata resin, astilbin from Engelhardtia roxburghiana (Huangqi) leaves, xanthohumol from hop (Humulus lupulus) extract, and cinnamaldehyde from cinnamon (Cinnamomum cassia) bark mixture(the *composition and proportional formulation of Inflamate® are detailed in Supplementary Table S2)*; Kenbics, Japan) was initially dissolved in an appropriate volume of dimethyl sulfoxide (DMSO; FUJIFILM Wako Pure Chemical Corporation, Osaka, Japan) to solubilize its bioactive components. The solution was subsequently centrifuged at 15,000 × g for 10 min, and the resulting supernatant was collected and stored at −80 °C until use. The stock concentration of Inflamate® was 3 mg/3 mL (1 mg/mL). Working solutions at various concentrations were prepared by serial two-fold dilution, and each was added to adherent cells at a ratio of 1:1000 (v/v, stock solution to culture medium) to determine the optimal treatment concentration for subsequent experiments.

#### 2.2.2. Cell Culture Medium for Experimental Treatments

The culture medium used for all experimental treatments consisted of high-glucose Dulbecco’s Modified Eagle Medium (DMEM) supplemented with 1% heat-inactivated fetal bovine serum (FBS) and 1% penicillin–streptomycin (P/S).

#### 2.2.3. Experimental Groups

Cells were divided into four experimental groups as follows (Figure S1): DMSO group: Cells were treated with DMSO vehicle alone for 48 h.

DMSO + LPS group: Cells were pre-treated with DMSO vehicle for 24 h, followed by stimulation with 1 μg/mL lipopolysaccharide (LPS) for an additional 24 h.

Inflamate® group: Cells were treated with Inflamate® alone for 24 h.

Inflamate® + LPS group: Cells were pre-treated with Inflamate® for 24 h, followed by stimulation with 1 μg/mL LPS for an additional 24 h.

Upon completion of each treatment, total RNA was extracted and gene expression was analyzed by qRT-PCR, as described in Section 2.4.

### 2.3. Cell Damage Assessment

Cell viability and cytotoxicity were evaluated using the CCK-8 and LDH assay kits (both from Dojindo Laboratories, Kumamoto, Japan), respectively, and compared with the control group to assess the extent of cellular damage. Cells from each experimental group were seeded in a 96-well plate and allowed to adhere for 24 h prior to treatment. After completion of the 48h treatment protocol described in Section 2.2.3., data for both CCK-8 and LDH assays were collected, and absorbance was measured at 450 nm (OD₄₅₀) using a microplate reader.

#### 2.3.1. CCK-8 Cell Viability Assay

Following completion of the treatment protocol, 100 μL of CCK-8 reagent solution was added to the culture medium at a ratio of 1:10 (reagent to medium). The cells were then incubated at 37 °C in the dark for 1 h. Subsequently, 100 μL aliquots were transferred to a new transparent 96-well plate (TPP Techno Plastic Products AG, Trasadingen, Switzerland), and the absorbance was measured at 450 nm (OD₄₅₀) using a microplate reader.

#### 2.3.2. LDH Cytotoxicity Assay

After the 48 h treatment period, the cell culture supernatant was collected from each well. A volume of 50 μL of the supernatant was transferred to a new 96-well plate (TPP Techno Plastic Products AG), followed by the addition of 50 μL of LDH detection reagent (1:1 ratio). The plate was incubated at room temperature in the dark for 30 min, and the absorbance was subsequently measured at 450 nm (OD₄₅₀).

### 2.4. Real-Time Quantitative Polymerase Chain Reaction (qPCR)

Total RNA was isolated from astrocyte cells treated with specific pharmacological agents using the FastGene RNA Basic Kit (Nippon Genetics Co., Ltd., Tokyo, Japan) in accordance with the manufacturer’s instructions. The concentration and purity of the extracted RNA were assessed using a NanoDrop 2000 spectrophotometer (Thermo Fisher Scientific, Waltham, MA, USA), and only samples with an A₂₆₀/A₂₈₀ ratio between 1.8 and 2.0 were used for subsequent analysis.

Complementary DNA (cDNA) was synthesized from 200 ng of total RNA using the GeneAce cDNA Synthesis Kit (NIPPON GENE Co., Ltd., Tokyo, Japan) in a 20 μL reaction volume, following the manufacturer’s protocol. Quantitative real-time PCR (qRT-PCR) was performed using the StepOnePlus Real-Time PCR System (Thermo Fisher Scientific) with SYBR™ Green qPCR Master Mix (Thermo Fisher Scientific). Each reaction was carried out in triplicate in a final volume of 10 μL, containing 5 μL of SYBR™ Green qPCR Master Mix (2×), 0.25 μL of Yellow Sample Buffer (40×), 0.5 μL each of forward and reverse primers (10 μM), 1 μL of cDNA template (10 ng/μL), and 2.75 μL of nuclease-free water.

Relative gene expression levels were quantified using the 2^−ΔΔ*CT*^ method, with ribosomal protein S13 (Rps13) serving as the internal reference gene. All procedures followed the manufacturer’s protocols and adhered to the Minimum Information for Publication of Quantitative Real-Time PCR Experiments (MIQE) guidelines[66]. The primer sequences used in this study are listed in Supplementary Table S1.

### 2.5. Statistical Analysis

All statistical analyses were performed using GraphPad Prism (version 11; GraphPad Software, San Diego, CA, USA) and Microsoft Excel for Microsoft 365 (Microsoft Corporation, Redmond, WA, USA). Data are presented as mean ± standard deviation (SD).

For cell viability (CCK-8) and cytotoxicity (LDH) assays, differences among groups were evaluated using one-way analysis of variance (ANOVA), followed by Dunnett’s post hoc multiple comparison tests, as appropriate.

For qRT-PCR gene expression data, two-way analysis of variance (ANOVA) was employed to evaluate the main effects of LPS stimulation and Inflamate® treatment, as well as their interaction. When significant effects were detected, Šídák’s multiple comparisons test was applied for post hoc analysis.

A p-value of < 0.05 was considered statistically significant. All experiments were performed independently with three biological replicates per condition.

## 3. Results

### 3.1. Determination of Non-Cytotoxic Concentrations of Inflamate® in Astrocytes

To establish the appropriate working concentration of Inflamate® for subsequent experiments, a dose–response assessment was performed using morphological observation, CCK-8 cell viability assay, and LDH cytotoxicity assay. Astrocytes were treated with Inflamate® at concentrations ranging from 0 to 0.3 μg/mL for 48 h, and cellular responses were evaluated.

#### 3.1.1. Morphological Assessment of Dose-Dependent Inflamate® Treatment

Phase-contrast microscopy revealed dose-dependent morphological changes in astrocytes following 48 h of Inflamate® treatment (Figure 1). At 0 μg/mL (untreated control), astrocytes exhibited a typical stellate morphology with well-spread processes and high cell density. At 0.0375 μg/mL, a moderate reduction in cell number was observed; however, the majority of cells retained their characteristic astrocytic morphology with extended processes, indicating that this concentration did not induce overt cytotoxicity. At 0.075 μg/mL, a further decrease in cell density was observed, accompanied by morphological alterations, including process retraction and partial cell-body rounding, suggesting the onset of cytotoxic effects. At 0.3 μg/mL, extensive cell death was evident, with most remaining cells displaying a rounded, detached morphology indicative of severe cytotoxicity (Figure 1).

**Figure 1.**
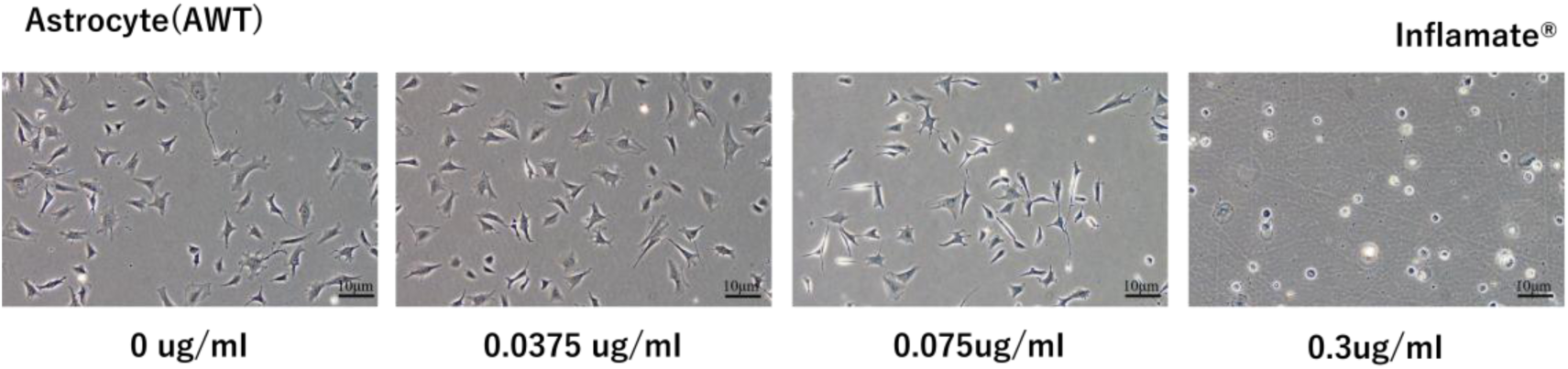
Dose-dependent morphological changes in mouse astrocytes (AWT) treated with Inflamate®. Representative phase-contrast micrographs of astrocytes following 48 h treatment with Inflamate® at concentrations of 0, 0.0375, 0.075, and 0.3 μg/mL. At 0 μg/mL, cells displayed a typical stellate morphology with well-spread processes. At 0.0375 μg/mL, cell morphology was largely preserved despite a moderate reduction in cell density. At 0.075 μg/mL, process retraction and cell body rounding were observed. At 0.3 μg/mL, extensive cell death with rounded, detached cells was evident. Scale bar information as indicated.

#### 3.1.2. CCK-8 Cell Viability Assay

Quantitative assessment of cell viability was performed using the CCK-8 assay (Figure 2a). Cell viability remained stable at lower concentrations of Inflamate®. No significant differences in absorbance (OD₄₅₀) were observed between the control group (0 μg/mL) and cells treated with 0.01875 μg/mL (p > 0.05, ns) or 0.0375 μg/mL (p > 0.05, ns). At 0.075 μg/mL, a modest but significant reduction in cell viability was detected (p < 0.01), consistent with the morphological changes observed at this concentration. At higher concentrations, cell viability was markedly decreased, with 0.15 μg/mL and 0.3 μg/mL both showing near-complete loss of metabolic activity (p < 0.0001), indicating severe cytotoxicity.

**Figure 2.**
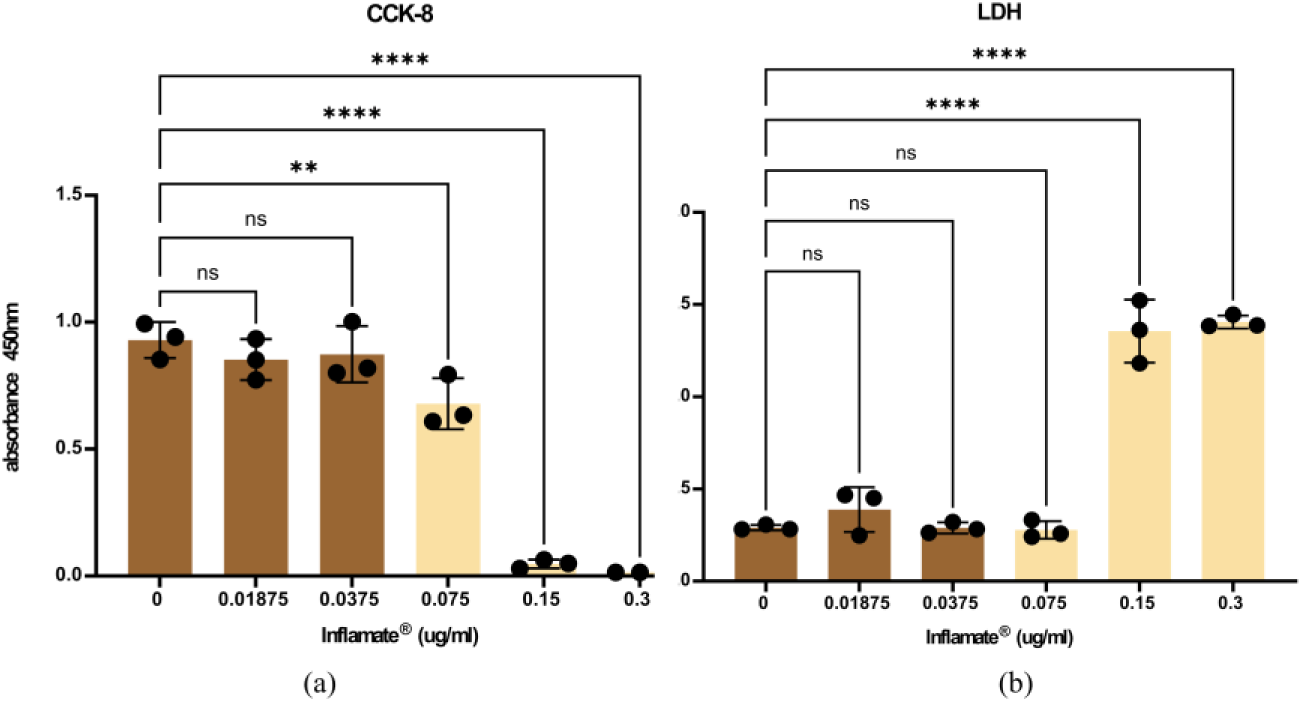
Cell viability and cytotoxicity assessment of Inflamate®-treated mouse astrocytes (AWT). (a) CCK-8 cell viability assay. Absorbance at 450 nm (OD₄₅₀) was measured following 48 h treatment with Inflamate® at concentrations of 0, 0.01875, 0.0375, 0.075, 0.15, and 0.3 μg/mL. No significant reduction in viability was observed at 0.01875 or 0.0375 μg/mL. A significant decrease was detected at 0.075 μg/mL (✱✱, p < 0.01), and near-complete loss of viability at 0.15 and 0.3 μg/mL (✱✱✱✱, p < 0.0001). (b) LDH cytotoxicity assay. LDH release remained at baseline levels at concentrations ≤ 0.075 μg/mL (ns for all). A significant increase in LDH release was observed at 0.15 and 0.3 μg/mL (✱✱✱✱, p < 0.0001). Data are presented as mean ± SD (n = 3–4 biological replicates per condition). Statistical analysis was performed using one-way ANOVA followed by Dunnett’s multiple comparisons test against the control group (0 μg/mL).

#### 3.1.3. LDH Cytotoxicity Assay

Cell membrane integrity was assessed using the LDH release assay as a complementary measure of cytotoxicity (Figure 2b). LDH release remained at baseline levels at concentrations of 0.01875, 0.0375, and 0.075 μg/mL, with no significant differences compared to the control group (p > 0.05, ns for all three concentrations). In contrast, a marked and significant increase in LDH release was observed at 0.15 μg/mL and 0.3 μg/mL (p < 0.0001 for both), confirming substantial membrane damage and cell death at these concentrations. Notably, while the CCK-8 assay detected a significant reduction in metabolic activity at 0.075 μg/mL, the LDH assay did not reveal significant membrane damage at this concentration, suggesting that the reduced viability at 0.075 μg/mL may reflect impaired cellular metabolism rather than overt cell lysis.

#### 3.1.4. Selection of Working Concentration

The results from morphological observation, CCK-8 cell viability assay, and LDH cytotoxicity assay are summarized in Table 1. Based on the integrated findings, 0.0375 μg/mL was selected as the optimal working concentration of Inflamate® for all subsequent experiments. At this concentration, astrocytes maintained normal stellate morphology, showed no significant reduction in cell viability (CCK-8, p > 0.05), and exhibited no detectable increase in membrane damage (LDH, p > 0.05), while remaining within a biologically active range. The next higher concentration (0.075 μg/mL) was excluded due to the significant reduction in CCK-8 viability (p < 0.01) and visible morphological alterations, despite the absence of significant LDH release.

**Table 1.** Summary of morphological observation, CCK-8 cell viability, and LDH cytotoxicity results for Inflamate®-treated astrocytes.

| Concentration ($\mu\text{g/mL}$) | Cell Morphology | CCK-8 Viability | LDH Cytotoxicity |
| --- | --- | --- | --- |
| 0 (Control) | Normal stellate | Baseline | Baseline |
| 0.01875 | Normal | ns | ns |
| **0.0375** ♦ | Normal (slight density ↓) | ns | ns |
| 0.075 | Process retraction, rounding | ** ($p < 0.01$) | ns |
| 0.15 | Severe damage | **** ($p < 0.0001$) | **** ($p < 0.0001$) |
| 0.3 | Rounded/detached (dead) | **** ($p < 0.0001$) | **** ($p < 0.0001$) |

<sup>1</sup> ns: not significant compared to control (Dunnett's test). ♦ Selected working concentration.

### 3.2. Morphological Changes in LPS-Stimulated Astrocytes Treated with Inflamate®

Preliminary morphological observations by phase-contrast microscopy suggested that LPS stimulation induced morphological changes consistent with reactive astrogliosis, and that Inflamate® treatment partially attenuated these changes (Supplementary Figure S1). However, given the limitations of phase-contrast imaging at this magnification, these observations are considered preliminary and require further validation using higher-magnification imaging or immunofluorescence staining for GFAP.

### 3.3. Inflamate® Selectively Suppresses LPS-Induced Pro-Inflammatory Gene Expression in Astrocytes

To investigate the anti-inflammatory effects of Inflamate® on astrocytes, we examined the expression of key inflammatory mediators by qRT-PCR (Figure 3). Under normal culture conditions (without LPS stimulation), Inflamate® treatment did not significantly alter the expression of any inflammatory gene examined compared to the DMSO vehicle control (p > 0.05 for all targets), indicating that the compound does not perturb baseline astrocyte homeostasis.

**Figure 3.**
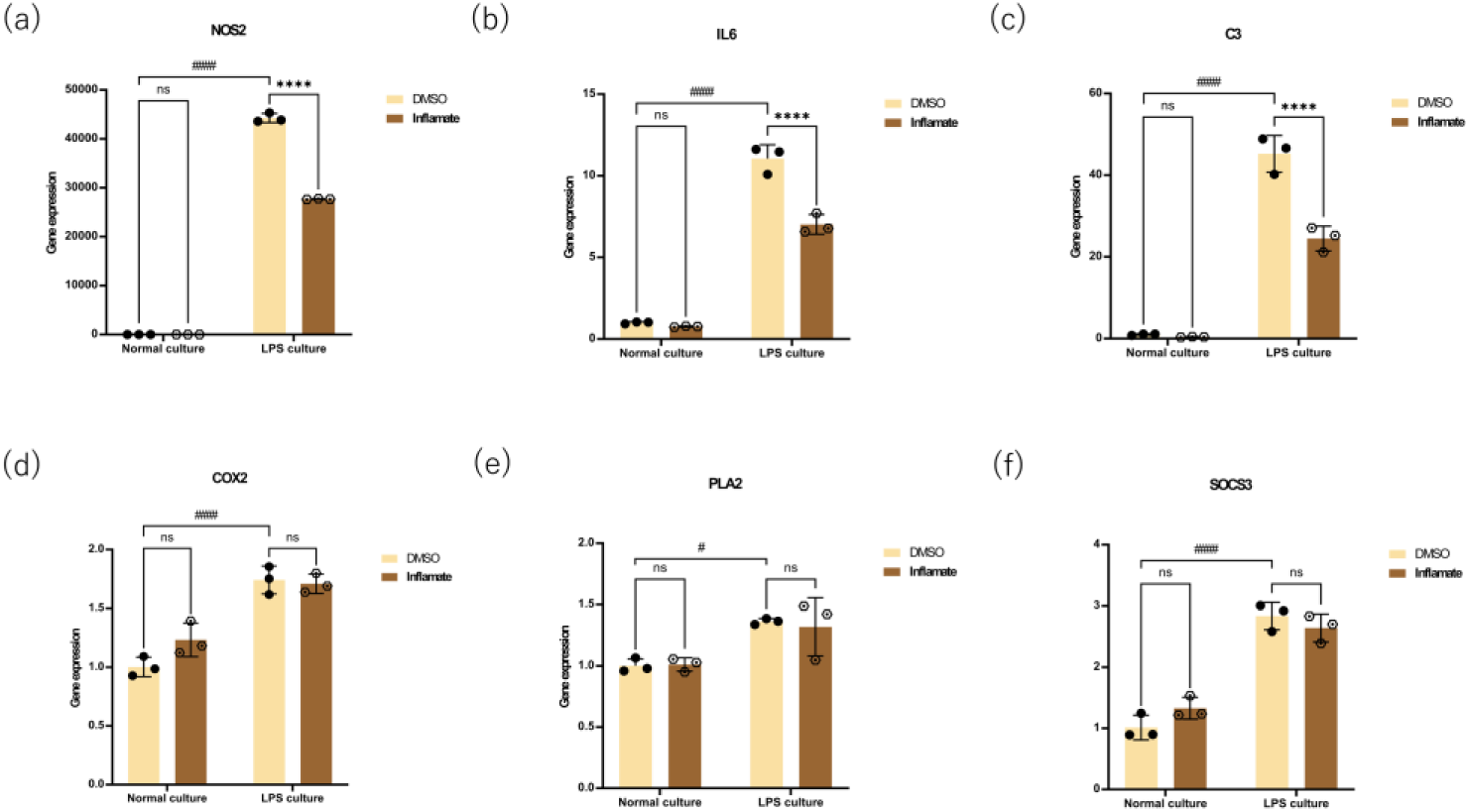
Inflamate® selectively suppresses LPS-induced pro-inflammatory gene expression in astrocytes without affecting the COX-2/prostaglandin pathway. Mouse astrocytes were treated with Inflamate® or DMSO vehicle control under normal culture conditions or following LPS stimulation (1 μg/mL) for 24 hours. Gene expression was quantified by qRT-PCR and normalized to the housekeeping gene. Data are presented as mean ± SD (n = 3 biological replicates per group). Statistical significance was determined by two-way ANOVA with appropriate post hoc test. ****p < 0.0001; ns, not significant. **(a)** NOS2 (iNOS) expression. Inflamate® significantly reduced LPS-induced NOS2 expression by approximately 64% compared to the DMSO control (p < 0.0001), with no effect under normal culture conditions. **(b)** IL6 expression. LPS-induced IL-6 was significantly suppressed by Inflamate® treatment (approximately 37% reduction; p < 0.0001), while baseline expression remained unaffected. **(c)** C3 (complement component 3) expression. LPS induced a dramatic upregulation of C3 (∼45-fold over normal culture) in DMSO-treated cells, which was significantly attenuated by Inflamate® to approximately 24-fold, representing an approximately 47% reduction relative to the LPS+DMSO control (p < 0.0001). **(d)** COX2 (cyclooxygenase-2) expression. LPS stimulation significantly induced COX2 expression compared to normal culture conditions (p < 0.0001). No significant differences were observed between Inflamate®- and DMSO-treated groups under either condition (p > 0.05), indicating that Inflamate® does not affect COX2 expression regardless of inflammatory context. **(e)** PLA2 *(PLA2g15, group XV phospholipase A2)* expression. Although LPS stimulation induced a modest but statistically significant increase in PLA2g15 expression (p < 0.05; Figure 3e), Inflamate® did not significantly alter PLA2g15 expression under either normal or LPS-stimulated conditions (p > 0.05), consistent with the compound not perturbing lysosomal phospholipid metabolic pathways. **(f)** SOCS3 expression. LPS stimulation significantly induced SOCS3 expression compared to normal culture conditions (p < 0.0001). No significant change was detected between Inflamate®- and DMSO-treated groups under either condition (p > 0.05).

LPS stimulation significantly induced the expression of NOS2, IL6, and C3 compared to normal culture conditions (p < 0.0001 for all three targets), confirming robust activation of the pro-inflammatory program. Although COX2, PLA2g15, and SOCS3 were all significantly induced by LPS (p < 0.0001, p < 0.05, and p < 0.0001, respectively; Figure 3d–3f), Inflamate® did not significantly alter the expression of any of these three genes compared to DMSO under either condition (p > 0.05). Upon LPS stimulation, DMSO-treated astrocytes exhibited robust upregulation of pro-inflammatory genes. Notably, Inflamate® treatment markedly suppressed LPS-induced *NOS2* (iNOS) expression to approximately 36% of the DMSO control level, representing an approximate 64% reduction (p < 0.0001; Figure 3a).

Similarly, *IL6* expression was significantly reduced by approximately 37% compared to the LPS + DMSO group (p < 0.0001; Figure 3b). *C3*, a complement component and established marker of neurotoxic A1 reactive astrocyte[37], was decreased by approximately 47% relative to the LPS+DMSO control (p < 0.0001; Figure 3c). The magnitude of C3 suppression is particularly noteworthy given that LPS induced a dramatic upregulation of C3 expression (approximately 45-fold relative to normal culture), and Inflamate® reduced this to approximately 24-fold, indicating a substantial attenuation of the A1 neurotoxic program.

In contrast, the expression of *COX2* (cyclooxygenase-2) and *PLA2* (PLA2g15, group XV phospholipase A2) was not significantly affected by Inflamate® treatment under either normal or LPS-stimulated conditions (p > 0.05; Figure 3d, 3e). Likewise, *SOCS3*, a negative feedback regulator of the JAK-STAT3 pathway, showed no significant change between Inflamate®- and DMSO-treated groups in either condition (p > 0.05; Figure 3f). These findings demonstrate selective suppression of the iNOS–NO and IL-6 signaling axes by Inflamate®, without affecting the COX-2/prostaglandin pathway. The lack of significant change in SOCS3 expression despite IL-6 suppression suggests that LPS-induced SOCS3 upregulation may be maintained through IL-6-independent mechanisms, as discussed in Section 4.1.

### 3.4. Inflamate® Attenuates Inflammation-Driven Iron Metabolism Dysregulation

Given the intimate crosstalk between neuroinflammation and iron dysregulation[28, 67], we next assessed the effect of Inflamate® on iron metabolism-related genes (Figure 4). Under normal culture conditions, no significant differences were observed between Inflamate®- and DMSO-treated astrocytes for any iron-related transcript examined (p > 0.05 for all targets).

**Figure 4.**
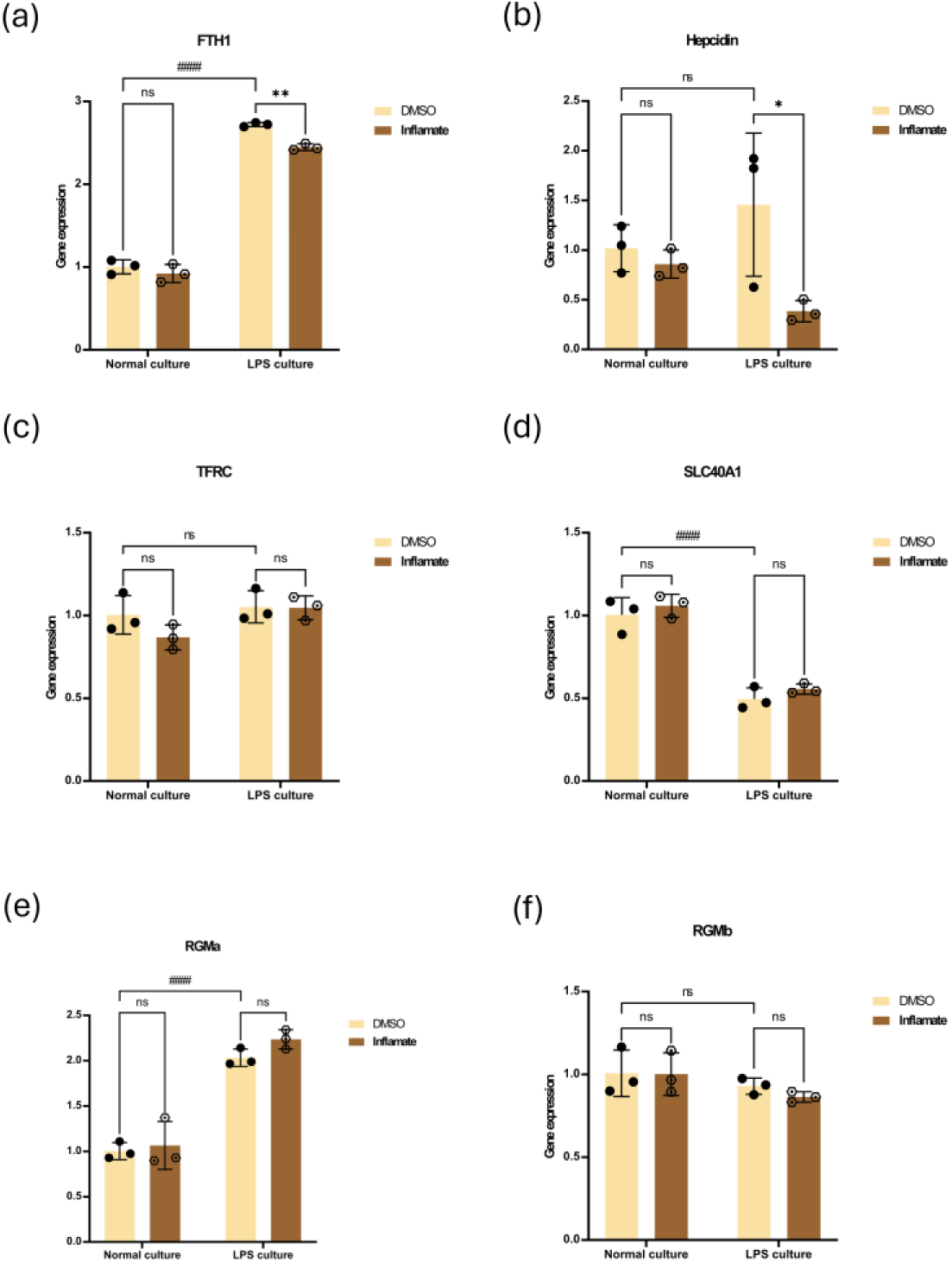
Inflamate® selectively attenuates inflammation-driven iron storage and hepcidin responses without disrupting iron transport machinery or BMP-SMAD signaling. Mouse astrocytes were treated with Inflamate® or DMSO vehicle control under normal culture conditions or following LPS stimulation (1 μg/mL) for 24 hours. Gene expression was quantified by qRT-PCR and normalized to the housekeeping gene. Data are presented as mean ± SD (n = 3 biological replicates per group). Statistical significance was determined by two-way ANOVA with appropriate post hoc test. *p < 0.05, **p < 0.01; ns, not significant. (a) FTH1 (ferritin heavy chain 1) expression. LPS stimulation induced FTH1 upregulation (∼2.7-fold) in DMSO-treated cells. Inflamate® significantly attenuated this response to approximately 2.5-fold (p < 0.01), with no effect under normal culture conditions. (b) Hepcidin (HAMP) expression. Inflamate® markedly reduced LPS-induced hepcidin expression to approximately 17% of the DMSO control level (p < 0.05), while baseline expression was unaffected. (c) TFRC (transferrin receptor 1) expression. LPS stimulation did not significantly alter TFRC expression compared to normal culture conditions (p > 0.05). No significant differences were observed between Inflamate®- and DMSO-treated groups under either condition (p > 0.05), indicating that iron uptake machinery remains intact. (d) SLC40A1 (ferroportin) expression LPS stimulation significantly reduced SLC40A1 expression compared to normal culture conditions (p < 0.0001), suggesting that acute TLR4 activation suppresses ferroportin-mediated iron export. No significant differences were observed between Inflamate®- and DMSO-treated groups under either condition (p > 0.05), indicating that Inflamate® does not further compromise ferroportin-mediated iron export. (e) RGMa expression. LPS stimulation significantly induced RGMa expression compared to normal culture conditions (p < 0.0001); however, no significant difference was observed between Inflamate®- and DMSO-treated groups under either condition (p > 0.05), indicating that Inflamate® does not interfere with LPS-induced BMP co-receptor upregulation. (f) RGMb expression. LPS stimulation did not significantly alter RGMb expression compared to normal culture conditions (p > 0.05). Expression was not significantly affected by Inflamate® in either normal or LPS conditions (p > 0.05), further confirming that the BMP-SMAD pathway remains intact and is not a target of Inflamate® action.

Under LPS-stimulated conditions, Inflamate® significantly reduced the expression of *FTH1* (ferritin heavy chain 1) compared to the DMSO control (p < 0.01; Figure 4a). LPS stimulation induced a marked upregulation of FTH1 in DMSO-treated cells (approximately 2.7-fold relative to normal culture), consistent with inflammation-driven intracellular iron sequestration. Inflamate® treatment attenuated this response to approximately 2.5-fold, representing a significant but moderate reduction. Furthermore,

LPS stimulation induced a significant upregulation of hepcidin expression compared to normal culture conditions (p < 0.05; Figure 4b), consistent with IL-6–STAT3-mediated HAMP transcriptional activation. Hepcidin (*HAMP*), the master regulator of iron homeostasis, was subsequently significantly downregulated by Inflamate® treatment under LPS conditions (p < 0.05; Figure 4b). Notably, hepcidin expression in the LPS + Inflamate® group was reduced to approximately 17% of the LPS + DMSO level, suggesting a substantial attenuation of the inflammation-driven hepcidin induction.

The concurrent reduction of both FTH1 and Hepcidin under LPS conditions is consistent with an overall attenuation of the inflammatory response rather than a direct suppression of iron storage capacity. Under inflammatory conditions, hepcidin promotes ferroportin degradation and consequent intracellular iron retention, which in turn drives compensatory FTH1 upregulation to safely sequester accumulated labile iron[19, 20]. The reduction of FTH1 by Inflamate® therefore likely reflects diminished upstream inflammatory signaling (NOS2, IL-6) and consequently reduced hepcidin-mediated iron-sequestering demand, rather than impairment of the intrinsic cytoprotective iron-buffering function of ferritin.

Importantly, the expression of *TFRC* (transferrin receptor 1, mediating iron uptake) and *SLC40A1* (ferroportin, mediating iron export) remained unchanged between Inflamate®- and DMSO-treated groups under both normal and LPS conditions (p > 0.05; Figure 4c, 4d). However, LPS stimulation per se significantly reduced SLC40A1 expression compared to normal culture condition(p < 0.0001; Figure 4d) suggesting that acute TLR4 activation suppresses ferroportin-mediated iron export independently of Inflamate® treatment. Similarly, *RGMa* and *RGMb*, which function as BMP co-receptors involved in hepcidin transcriptional regulation via the BMP-SMAD pathway[68, 69], were not significantly altered by Inflamate® treatment (p > 0.05; Figure 4e, 4f). LPS stimulation significantly induced RGMa expression compared to normal culture conditions (p < 0.0001; Figure 4e), whereas Inflamate® did not significantly alter RGMa expression under either condition (p > 0.05), suggesting that Inflamate® does not interfere with LPS-induced BMP co-receptor upregulation. These results indicate that Inflamate® does not broadly disrupt iron transport machinery or BMP-SMAD signaling, but rather selectively attenuates the inflammation-driven (IL-6–STAT3-mediated) upregulation of hepcidin and its downstream iron storage response. Notably, LPS stimulation per se did not significantly alter the expression of TFRC or RGMb compared to normal culture conditions (p > 0.05 for both), suggesting that acute TLR4 activation does not substantially perturb all components of the core iron transport machinery or BMP-SMAD signaling under the present experimental conditions.

### 3.5. Inflamate® Promotes a Shift Toward the Neuroprotective A2 Astrocyte Phenotype

To assess whether Inflamate® influences astrocyte reactive polarization, we examined markers associated with neurotoxic A1 and neuroprotective A2 astrocyte phenotypes (Figure 5). As described in Section 3.3, the A1 marker *C3* was significantly downregulated by Inflamate® under LPS conditions (p < 0.0001). Conversely, *S100A10*, an established marker of the neuroprotective A2 phenotype[43, 70], was significantly upregulated by Inflamate® in both normal culture conditions (p < 0.05; Figure 5b) and LPS-stimulated conditions (p < 0.01; Figure 5b). Notably, LPS stimulation alone did not significantly alter S100A10 expression compared to normal culture conditions (p > 0.05; Figure 5b), indicating that the Inflamate®-mediated upregulation of S100A10 is independent of the inflammatory context and likely reflects a constitutive pro-A2 transcriptional effect of the compound. S100A10 was the only gene among all 14 targets examined that showed a significant increase in expression upon Inflamate® treatment, further underscoring the specificity of this constitutive pro-A2 transcriptional effect among the full panel of targets assessed.

**Figure 5.**
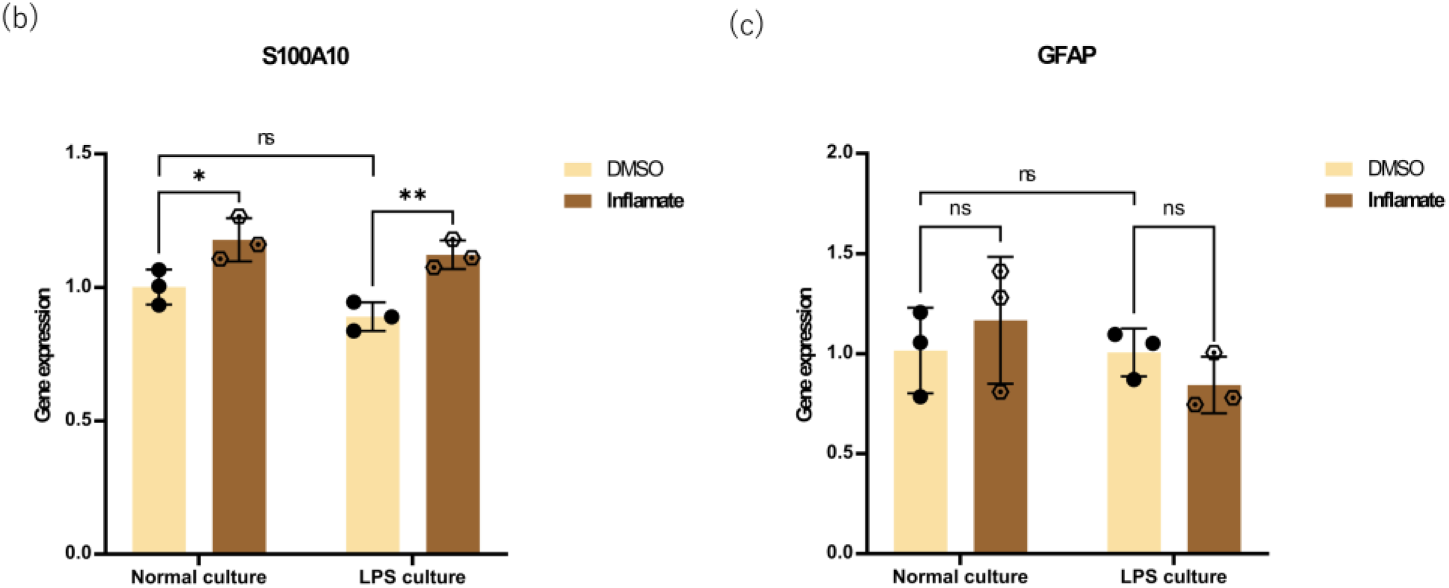
Inflamate® promotes a phenotypic shift from neurotoxic A1 toward neuroprotective A2 astrocyte polarization. Mouse astrocytes were treated with Inflamate® or DMSO vehicle control under normal culture conditions or following LPS stimulation (1 μg/mL) for 24 hours. Gene expression was quantified by qRT-PCR and normalized to the housekeeping gene. Data are presented as mean ± SD (n = 3 biological replicates per group). Statistical significance was determined by two-way ANOVA with appropriate post hoc test. *p < 0.05, **p < 0.01, ****p < 0.0001; ns, not significant. (a) C3 (complement component 3; A1 neurotoxic marker) expression. LPS induced a dramatic upregulation of C3 in DMSO-treated cells, which was significantly attenuated by Inflamate® (approximately 47% reduction relative to the LPS+DMSO control; p < 0.0001). No significant difference was observed under normal culture conditions. (as shown in Figure 3c) (b) S100A10 (A2 neuroprotective marker) expression. Inflamate® significantly upregulated S100A10 under both normal culture conditions (p < 0.05) and LPS-stimulated conditions (p < 0.01). S100A10 was the only gene among all 14 targets examined that showed a significant increase upon Inflamate® treatment regardless of inflammatory context, suggesting a constitutive pro-A2 effect. (c) GFAP (general astrocyte reactivity marker) expression. No significant differences were observed between Inflamate®-and DMSO-treated groups under either condition (p > 0.05), indicating that Inflamate® does not globally suppress astrocyte activation but rather selectively modulates the qualitative nature of the reactive response.

**Figure 6.**
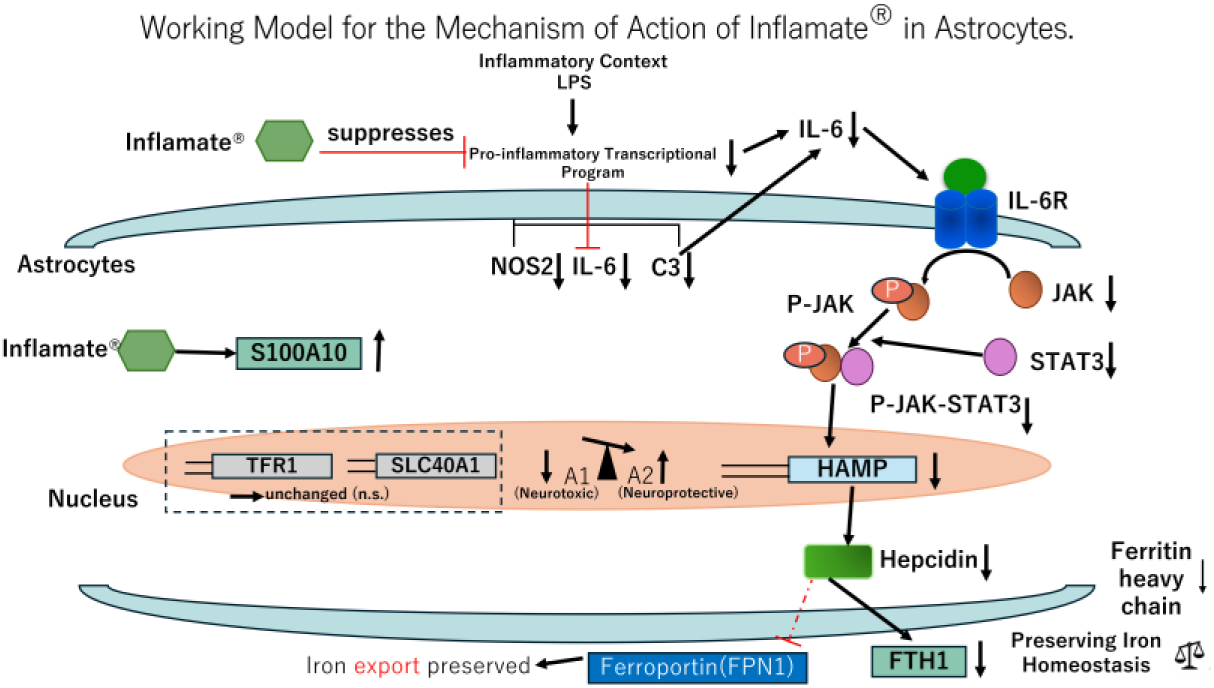
Working model for the mechanism of action of Inflamate® in astrocytes. Inflamate® suppresses LPS-induced pro-inflammatory transcriptional programs in astrocytes, reducing the expression of NOS2, IL-6, and C3. Decreased IL-6 attenuates the IL-6R–JAK–STAT3 signaling cascade, leading to reduced HAMP transcription, lower hepcidin levels, and attenuated FTH1 upregulation. Concurrently, Inflamate® upregulates S100A10 in a manner partially independent of the inflammatory context, while preserving basal iron transport mechanisms (TFRC and SLC40A1 remained unchanged), and shifting the astrocyte phenotype from neurotoxic A1 toward neuroprotective A2. Together, these actions reflect a coordinated, multi-target neuroprotective mechanism relevant to neuroinflammation and iron dysregulation in neurodegenerative disease. ↓, downregulation; ↑, upregulation; ⊣, inhibition.

The expression of *GFAP*, a general marker of astrocyte reactivity, was not significantly altered by Inflamate® treatment under either condition (p > 0.05; Figure 5c). LPS stimulation did not significantly alter GFAP expression compared to normal culture conditions (p > 0.05; Figure 5c), suggesting that the acute TLR4-mediated inflammatory stimulus employed in this study was insufficient to induce a detectable transcriptional upregulation of this general reactivity marker within the 24-hour observation window. This finding indicates that Inflamate® does not suppress astrocyte activation globally but rather selectively modulates the qualitative nature of the reactive response—shifting the phenotypic balance without abolishing reactivity itself.

Taken together, the concurrent downregulation of C3 (A1 marker) and upregulation of S100A10 (A2 marker) strongly suggest that Inflamate® promotes a phenotypic shift from the neurotoxic A1 toward the neuroprotective A2 astrocyte state. This pattern is consistent with recent reports demonstrating that anti-inflammatory interventions can simultaneously suppress C3 and iNOS while promoting S100A10 and Arg1 expression, thereby exerting neuroprotective effects[45, 50, 71]. The combined attenuation of pro-inflammatory mediators (NOS2, IL-6, C3), modulation of inflammation-driven iron metabolic programs (Hepcidin, FTH1), and promotion of the A2 neuroprotective phenotype (S100A10) points to a coordinated mechanism by which Inflamate® may exert neuroprotective effects in the context of neuroinflammation.

## 4. Discussion

Astrocytes are the most abundant glial cells in the central nervous system and play pivotal roles in maintaining neuronal homeostasis, modulating synaptic activity, and regulating the blood–brain barrier[5, 72]. However, under pathological conditions such as neurodegeneration, traumatic brain injury, and systemic infection, astrocytes undergo reactive astrogliosis—a heterogeneous response that can adopt either neurotoxic (A1) or neuroprotective (A2) phenotypes[37, 50, 73]. The neurotoxic A1 phenotype, characterized by upregulation of complement component C3 and pro-inflammatory mediators including iNOS and IL-6, has been implicated in neuronal and oligodendrocyte death in multiple neurodegenerative diseases[37, 74]. Conversely, the A2 phenotype, marked by S100A10 expression, promotes neuronal survival, synaptogenesis, and tissue repair[43, 44, 50]. Identifying compounds that can selectively suppress the A1 program while promoting the A2 phenotype therefore represents a promising therapeutic strategy for neuroinflammatory and neurodegenerative conditions.

In the present study, we demonstrate that Inflamate®, at a non-cytotoxic concentration (0.0375 μg/mL), selectively modulates LPS-induced astrocyte inflammatory responses, attenuates inflammation-driven iron metabolic dysregulation, and promotes a phenotypic shift toward the neuroprotective A2 state. These findings collectively suggest a coordinated, multi-target anti-neuroinflammatory mechanism that warrants further investigation.

### 4.1. Selective Anti-Inflammatory Profile of Inflamate®

A key finding of this study is the pathway-selective nature of the anti-inflammatory action of Inflamate®. The compound significantly suppressed LPS-induced NOS2 (approximately 64% reduction), IL-6 (approximately 37% reduction), and C3 (approximately 47% reduction), while leaving COX-2, PLA2g15, and SOCS3 expression unaffected (Figure 3). This pattern indicates that Inflamate® does not act as a broad-spectrum anti-inflammatory agent but rather targets specific signaling nodes within the inflammatory cascade.

The robust suppression of NOS2 and IL-6 suggests that Inflamate® may interfere with the NF-κB and/or JAK-STAT3 signaling pathways, which are the principal transcriptional regulators of these genes in LPS-stimulated astrocytes[75–77]. NF-κB activation is a central mediator of TLR4-dependent inflammatory signaling, driving the transcription of iNOS, IL-6, and numerous other pro-inflammatory mediators[78–80]. The concurrent reduction of both NOS2 and IL-6 is consistent with upstream inhibition at or above the level of NF-κB, although the precise molecular target remains to be elucidated.

In contrast, Inflamate® did not significantly alter the expression of Cox2 or Pla2g15 compared with DMSO under either condition (p > 0.05; Figure 3d,e), despite their significant induction by LPS (p < 0.0001 and p < 0.05, respectively). Notably, the LPS-induced upregulation of Cox2 was of modest magnitude (∼1.5-fold) relative to the dramatic induction of Nos2 and C3 under identical conditions, suggesting that the biological contribution of COX-2 to the acute neuroinflammatory response in this astrocyte model may be relatively limited. Nevertheless, the lack of effect on COX-2 is mechanistically informative, as COX-2 represents a parallel but distinct arm of the inflammatory response—the arachidonic acid/prostaglandin pathway [81–83]. From a therapeutic perspective, this selectivity may be advantageous, since COX-2-derived prostaglandins also play homeostatic and resolution-phase roles in neuroinflammation [84], and their indiscriminate suppression could impede tissue repair processes. The parallel lack of effect on Pla2g15 (group XV phospholipase A2)—a lysosomal phospholipase involved in phospholipid catabolism rather than the classical inflammatory arachidonic acid cascade [85], urther indicates that Inflamate® does not perturb general phospholipid metabolic pathways. Taken together, these observations support the view that Inflamate® selectively targets the NF-κB/iNOS and JAK–STAT3 axes while sparing the COX-2/prostaglandin and phospholipid metabolic pathways.

The unchanged expression of SOCS3, a canonical negative feedback regulator of the JAK-STAT3 pathway[86], may appear paradoxical given the significant reduction in IL-6. However, this observation is consistent with the interpretation that LPS-induced SOCS3 upregulation is maintained through IL-6-independent mechanisms—such as direct NF-κB-mediated transcription or inputs from other cytokines (e.g., IL-10, IFN-γ) that were not modulated under the present experimental conditions[87] rather than being solely driven by IL-6–JAK-STAT3 signaling. Under this interpretation, the suppression of IL-6 by Inflamate® would be insufficient to reduce SOCS3 expression, as the dominant driver of SOCS3 induction under LPS conditions may operate independently of the IL-6 axis. Time-course studies examining earlier time points (e.g., 4–12 hours) and pathway-specific inhibitor experiments would help delineate the relative contributions of IL-6-dependent and IL-6-independent mechanisms to SOCS3 regulation in LPS-stimulated astrocytes.

### 4.2. Attenuation of Inflammation-Driven Iron Dysregulation

The crosstalk between neuroinflammation and iron metabolism is increasingly recognized as a critical pathological axis in neurodegenerative diseases[23, 28]. Under inflammatory conditions, IL-6 activates the JAK-STAT3 pathway to induce hepcidin transcription, which in turn promotes ferroportin internalization and degradation, leading to intracellular iron retention[88, 89]. The accumulated labile iron catalyzes Fenton chemistry-driven oxidative stress, contributing to ferroptosis and neuronal damage[90, 91]. Compensatory upregulation of ferritin (FTH1) serves to safely sequester excess intracellular iron, but this protective mechanism may be overwhelmed under sustained inflammatory conditions[24].

Our findings demonstrate that Inflamate® significantly reduced both hepcidin (to approximately 17% of the LPS + DMSO level) and FTH1 expression under LPS-stimulated conditions (Figure 4a, 4b), while leaving the core iron transport machinery (TFRC, SLC40A1) and BMP-SMAD pathway components (RGMa, RGMb) unaffected (Figure 4c–4f). This pattern strongly suggests that the iron-modulatory effects of Inflamate® are secondary to its anti-inflammatory action rather than reflecting direct interference with iron homeostatic pathways. Specifically, the reduction of IL-6 by Inflamate® would be expected to diminish IL-6–STAT3-mediated hepcidin induction, thereby alleviating hepcidin-driven intracellular iron accumulation and reducing the compensatory demand for FTH1 upregulation. Although the reduction in FTH1 reached statistical significance, the modest magnitude of this effect (∼14%, from approximately 2.7-fold to 2.5-fold) suggests that the biological impact on iron storage capacity may be limited under the present experimental conditions. The primary iron-modulatory effect of Inflamate® is therefore more accurately reflected by the marked suppression of hepcidin, which represents the upstream regulatory node of the inflammation–iron retention axis.

The preservation of TFRC and SLC40A1 expression is particularly important from a safety perspective, as disruption of these transporters could lead to either iron deficiency (if uptake is impaired) or iron overload (if export is blocked)[20]. Similarly, the unchanged expression of RGMa and RGMb indicates that the BMP-SMAD pathway—the principal homeostatic regulator of hepcidin under non-inflammatory conditions[31, 68]—remains intact, further supporting the conclusion that Inflamate® selectively targets the inflammation-driven (IL-6–STAT3) arm of hepcidin regulation.

These findings have potential implications for neurodegenerative conditions in which inflammation-driven iron accumulation contributes to disease progression. In Alzheimer’s disease, Parkinson’s disease, and multiple sclerosis, astrocytic hepcidin upregulation and consequent iron dysregulation have been implicated in oxidative damage and neuronal loss[28, 91–93]. By attenuating this pathological axis, Inflamate® may help preserve iron homeostasis in the neuroinflammatory milieu. This mechanism may also be relevant in the context of exercise physiology, as strenuous physical activity has been shown to elevate systemic IL-6 levels and increase blood–brain barrier permeability[7], while gut-derived endotoxemia following exhaustive exercise may further introduce circulating LPS into the systemic circulation[94, 95], thereby propagating peripheral inflammatory signals into the CNS. To the extent that the IL-6–JAK-STAT3–hepcidin axis activated under LPS conditions shares mechanistic features with exercise-induced glial signaling, the present findings raise the possibility that hepcidin-mediated iron dysregulation may also occur in glial cells[54] under conditions of repeated exhaustive exertion—a hypothesis that warrants direct investigation in exercise-relevant models. Although peripheral and central cytokine compartments are not always tightly coupled during acute exercise [96], this possibility is particularly pertinent under conditions of repeated, exhaustive exertion, where sustained IL-6 elevation may be sufficient to engage central hepcidin regulation[97, 98]. Exercise-induced neuroinflammation and associated ferroptotic vulnerability therefore represent an emerging area of concern for athletes and physically active populations, and nutritional strategies capable of selectively attenuating the IL-6–hepcidin axis without disrupting basal iron transport may offer a targeted approach to preserving CNS iron homeostasis during and after intense physical exertion.

### 4.3. Promotion of the Neuroprotective A2 Astrocyte Phenotype

While the A1/A2 binary classification has been subject to ongoing refinement, and astrocyte reactive states are now understood to span a transcriptional continuum [103, 104], the directional modulation of established phenotypic markers—specifically C3 as an A1-associated marker and S100A10 as an A2-associated marker—provides a tractable framework for assessing the qualitative shift in astrocyte reactivity induced by Inflamate®. Perhaps the most striking finding of this study is the concurrent downregulation of the A1 marker C3 and upregulation of the A2 marker S100A10 by Inflamate® (Figure 5). This bidirectional modulation of astrocyte polarization markers suggests that Inflamate® does not merely suppress inflammation but actively promotes a phenotypic shift toward the neuroprotective A2 state.

The upregulation of S100A10 is particularly noteworthy for several reasons. First, S100A10 was the only gene among all 14 targets examined that showed a significant increase upon Inflamate® treatment, highlighting the specificity of this effect. Second, this upregulation was observed under both normal and LPS-stimulated conditions, suggesting a constitutive pro-A2 mechanism that is independent of the inflammatory context. This is distinct from the anti-inflammatory effects on NOS2, IL-6, and C3, which were only manifest under LPS stimulation. Third, S100A10 (also known as p11) has been independently implicated in neuroprotection, serotonin receptor trafficking, and antidepressant responses[99, 100], suggesting that its upregulation may confer benefits beyond the anti-inflammatory context.

The unchanged expression of GFAP further supports the interpretation that Inflamate® modulates the quality rather than the magnitude of astrocyte reactivity. GFAP upregulation is a hallmark of reactive astrogliosis regardless of the A1/A2 polarization state[43]. The preservation of GFAP expression indicates that Inflamate®-treated astrocytes remain capable of mounting a reactive response—which is essential for scar formation, blood–brain barrier repair, and containment of injury[101]—while being redirected toward a neuroprotective functional program.

The mechanism by which Inflamate® upregulates S100A10 remains to be determined. Possible pathways include activation of STAT6 signaling (which has been associated with A2 polarization)[102, 103], modulation of epigenetic regulators that control S100A10 transcription, or indirect effects mediated through the suppression of A1-promoting signals (e.g., NF-κB, C3) that may normally antagonize A2 gene expression[70]. Future studies employing transcriptomic profiling and pathway inhibitor experiments will be necessary to delineate the upstream regulators of this effect.

### 4.4. Integrated Mechanistic Model

Integrating the findings from all experimental endpoints, we propose the following working model for the mechanism of action of Inflamate® in astrocytes: Inflamate® suppresses LPS-induced activation of pro-inflammatory transcriptional programs, reducing the expression of NOS2, IL-6, and C3. The reduction of IL-6 at the transcriptional level is consistent with attenuated IL-6–STAT3- mediated hepcidin induction, which would be expected to reduce the downstream drive for ferroportin internalization, intracellular iron retention, and compensatory FTH1 upregulation. Concurrently, Inflamate® promotes S100A10 expression through a mechanism that is at least partially independent of the inflammatory context, shifting the astrocyte phenotypic balance from neurotoxic A1 toward neuroprotective A2. This coordinated modulation of inflammatory, iron metabolic, and astrocyte polarization gene programs constitutes a multi-target neuroprotective transcriptional signature that is particularly relevant to neurodegenerative conditions characterized by chronic neuroinflammation and iron dysregulation. Critically, these transcriptional effects are achieved without disrupting the expression of core iron transport genes (TFRC, SLC40A1) or BMP-SMAD pathway components (RGMa, RGMb), and without globally suppressing astrocyte reactivity (GFAP unchanged), indicating a targeted rather than broadly suppressive transcriptional mechanism.

Moreover, given that strenuous exercise can induce transient but significant neuroinflammatory responses—including elevated CNS IL-6 signaling[104] and reactive astrocyte polarization toward the neurotoxic A1 phenotype[51, 105]—if the overlap between LPS-induced and exercise-induced activation of the IL-6–JAK-STAT3–hepcidin axis is confirmed in future studies, the multi-target profile of Inflamate® may offer a relevant nutritional strategy for supporting neurological recovery and CNS iron homeostasis in physically active populations.

### 4.5. Limitations and Future Directions

Several limitations of the present study should be acknowledged. First, our experiments were conducted in mouse astrocyte monocultures, which do not fully recapitulate the complex multicellular interactions of the CNS microenvironment. The effects of Inflamate^®^ on astrocyte–neuron, astrocyte–microglia, and astrocyte–oligodendrocyte crosstalk remain to be investigated using co-culture systems or organotypic slice cultures. Second, LPS stimulation, while a well-established model of TLR4- mediated neuroinflammation, does not fully replicate the chronic, multifactorial inflammatory milieu present in neurodegenerative diseases. Validation using disease-relevant stimuli (e.g., amyloid-β, α-synuclein aggregates) and in vivo models will be essential. Third, the molecular targets of Inflamate® have not been identified in this study. Determining whether the compound directly inhibits NF-κB, modulates upstream kinases, or acts through alternative mechanisms will require targeted pathway analysis, reporter assays, and potentially unbiased approaches such as thermal proteome profiling or activity-based protein profiling. Fourth, the present study examined gene expression at a single time point (24 hours post-LPS stimulation). Time-course experiments and protein-level validation (e.g., Western blot and ELISA for secreted IL-6 and NO) would strengthen the conclusions and provide insight into the kinetics of Inflamate® action. Finally, as noted above, the A1/A2 classification employed in this study represents an operational simplification of a transcriptionally continuous spectrum of astrocyte reactive states[106, 107]. While this framework is sufficient to capture the directional phenotypic shift induced by Inflamate® at the level of key marker genes, single-cell transcriptomic approaches would provide a more comprehensive and unbiased characterization of the full spectrum of astrocyte heterogeneity affected by the compound.

Lastly, the present study employed LPS as a model of acute neuroinflammation, which, while mechanistically well-defined, does not directly replicate exercise-induced inflammatory conditions. However, the shared involvement of the IL-6–JAK-STAT3–hepcidin axis in both infection-driven and exercise-induced neuroinflammation[65] suggests that the mechanisms identified here may be translatable to exercise-related contexts. Future studies employing exercise-mimetic stimuli (e.g., recombinant IL-6 at physiologically relevant concentrations, conditioned media from exercised muscle cells) or in vivo exercise models would be valuable for determining whether Inflamate® can attenuate exercise-induced neuroinflammatory responses and preserve CNS iron homeostasis in physically active subjects. Such investigations would further strengthen the relevance of the present findings to the field of sports nutrition and exercise neuroscience.

## 5. Conclusions

In this study, we demonstrate that Inflamate®, at a non-cytotoxic concentration (0.0375 μg/mL), exerts selective anti-inflammatory, iron-modulatory, and astrocyte phenotype-shifting effects in LPS-stimulated mouse astrocytes at the transcriptional level. The principal findings are as follows:

Selective anti-inflammatory action: Inflamate® significantly suppressed LPS-induced expression of NOS2 (approximately 64% reduction), IL-6 (approximately 37% reduction), and C3 (approximately 47% reduction relative to the LPS+DMSO control), while COX-2, PLA2g15(a lysosomal phospholipase), and SOCS3 remained unaffected, demonstrating selective targeting of the iNOS–NO and IL-6 signaling axes at the transcriptional level rather than the COX-2/prostaglandin or phospholipid metabolic pathways.

Attenuation of inflammation-driven iron dysregulation: Inflamate® reduced LPS-induced hepcidin expression to approximately 17% of the control level and significantly attenuated FTH1 upregulation, without altering the expression of core iron transporters (TFRC, SLC40A1) or BMP-SMAD pathway components (RGMa, RGMb). These transcriptional findings are consistent with selective attenuation of the IL-6–STAT3–hepcidin axis as the primary iron-modulatory mechanism of Inflamate®, without broad disruption of iron homeostatic gene programs.

Promotion of the neuroprotective A2 phenotype: Inflamate® concurrently downregulated the A1 neurotoxic marker C3 and upregulated the A2 neuroprotective marker S100A10 under both basal and inflammatory conditions, while GFAP expression remained unchanged. This transcriptional pattern demonstrates a qualitative shift in astrocyte reactivity from neurotoxic toward neuroprotective without abolishing the reactive response itself.

Safety profile: At the selected working concentration, Inflamate® did not alter baseline gene expression of any target examined (except S100A10), did not reduce cell viability, and did not induce membrane damage, supporting a favorable safety profile for astrocyte applications.

Taken together, these results identify Inflamate® as a selective, multi-target modulator of neuroinflammation-associated transcriptional pathology, coordinating the suppression of pro-inflammatory gene expression, attenuation of inflammation-driven iron metabolic gene dysregulation, and promotion of the neuroprotective A2 astrocyte transcriptional phenotype. Notably, insofar as the IL-6–JAK-STAT3–hepcidin axis activated under LPS conditions shares mechanistic features with exercise-induced systemic inflammation, these findings may extend beyond neurodegenerative disease to exercise-related neuroinflammatory contexts, where targeted nutritional interventions capable of preserving CNS iron homeostasis and supporting neuroprotective glial function are of growing interest. Future studies employing in *vivo* neurodegenerative and exercise models, protein-level validation, and mechanistic pathway analyses are warranted to confirm these transcriptional findings and elucidate the precise molecular targets of Inflamate®.

## 6. Appendix A. Supplementary data

### Supplementary Materials

**Table S1:**
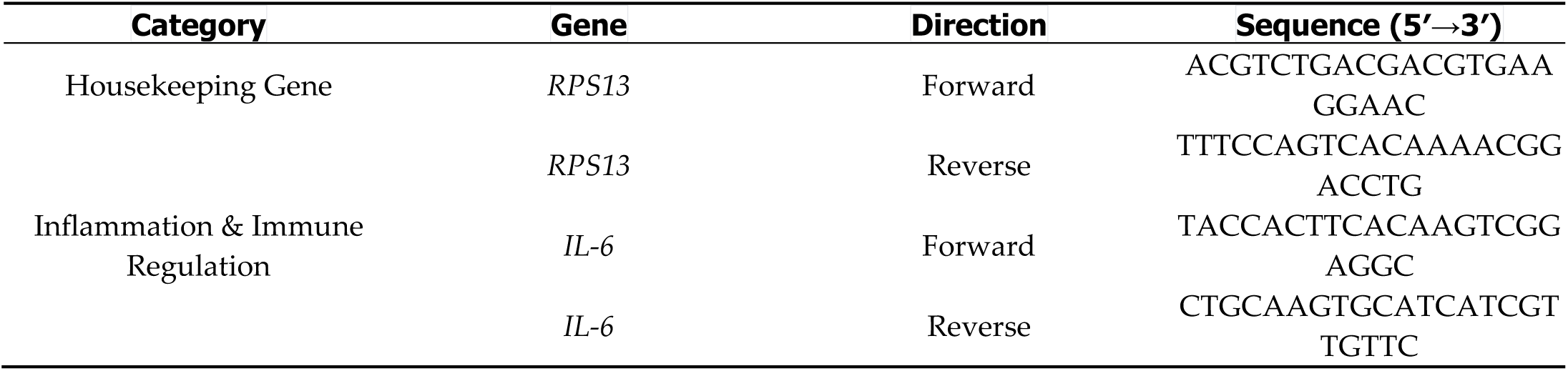
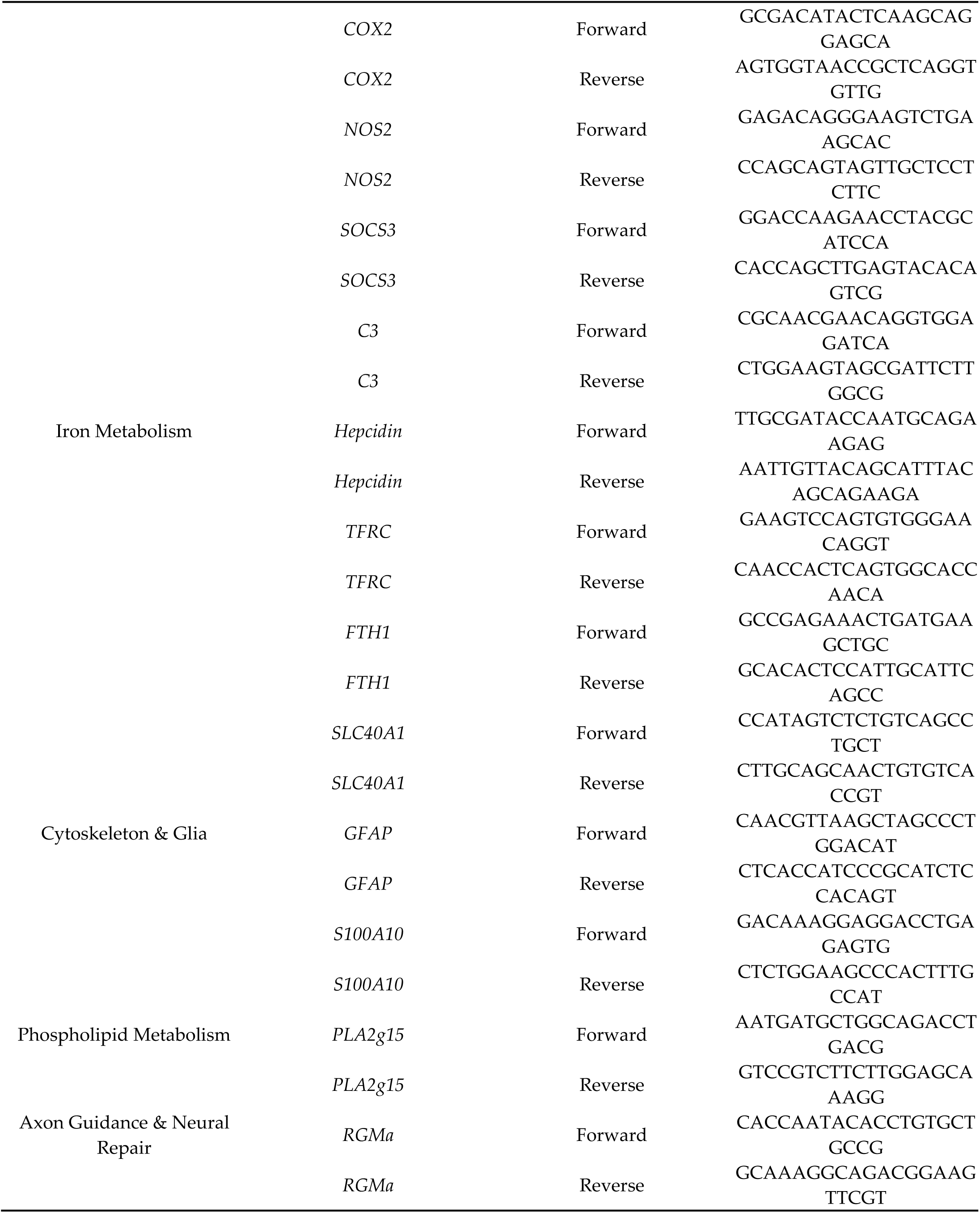
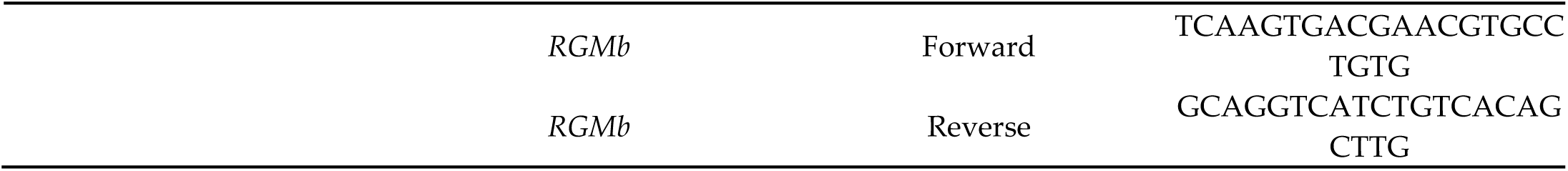
Primer sequences used for qRT-PCR analysis in this study.

**Table S2.** Composition and proportional formulation of Inflamate® dietary supplement. ¹ Rounding of individual component percentages results in a total slightly exceeding 100%. The remaining mass per capsule (approximately 149.7 mg) consists of excipients (capsule shell and other inactive ingredients) not disclosed by the manufacturer.

| Component | Botanical Source | Content per Capsule (mg) | Proportion (% w/w) |
| --- | --- | --- | --- |
| Boswellic acids | *Boswellia serrata* (resin extract) | 42.9 | 28.6 |
| Astilbin | *Engelhardtia roxburghiana* (Huangqi leaf extract) | 42.9 | 28.6 |
| Xanthohumol | *Humulus lupulus* (hop dry extract) | 42.9 | 28.6 |
| Cinnamaldehyde | *Cinnamomum cassia* (cinnamon bark powder) | 21.6 | 14.4 |
| **Total active ingredients** | — | **150.3** | **100.2 <sup>1</sup>** |

**Figure S1.**
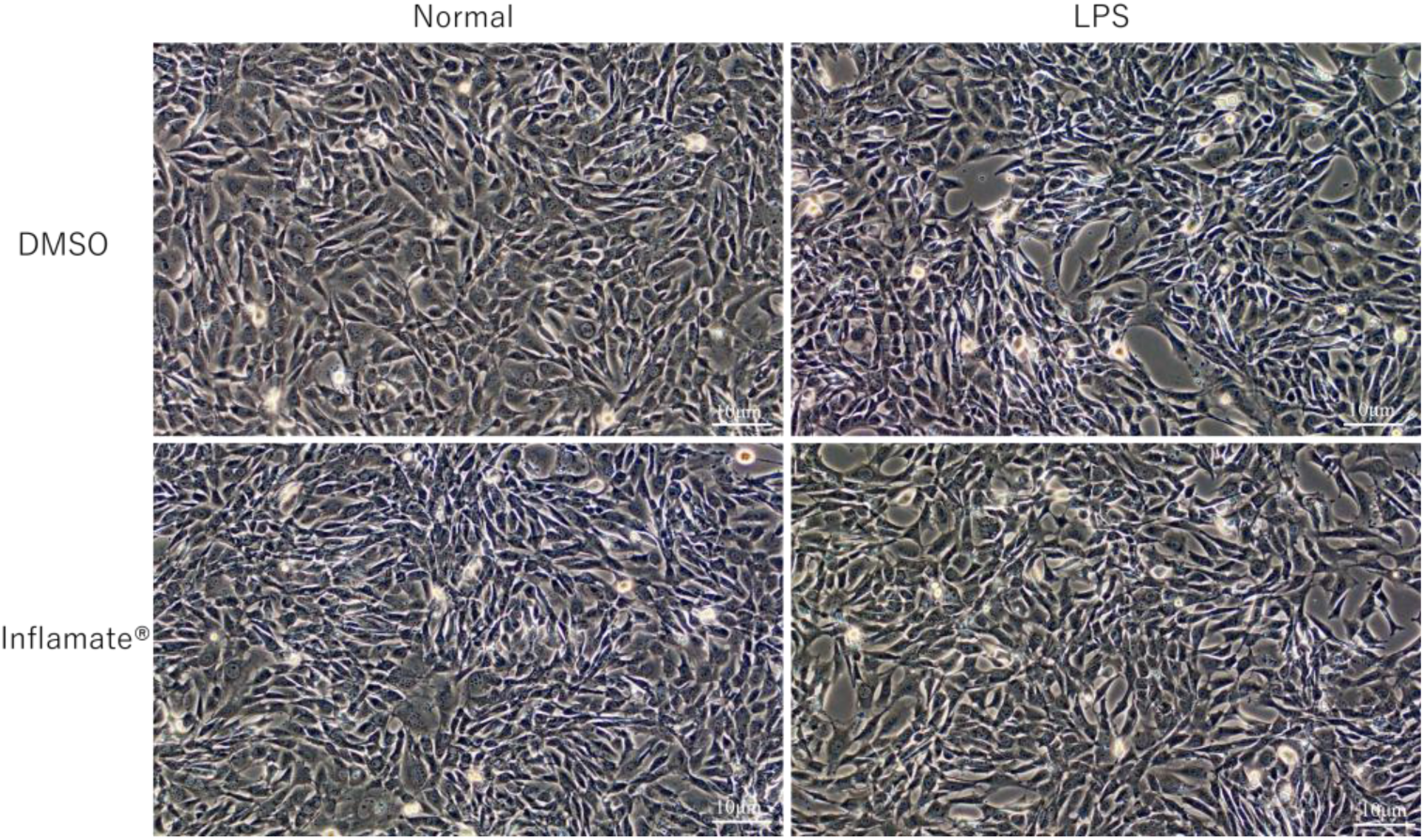
Morphological assessment of astrocytes (AWT) across four experimental groups. Representative phase-contrast micrographs of astrocytes following the 48 h treatment protocol. Top left: Normal + DMSO (vehicle control), showing typical stellate morphology with well-organized processes. Top right: LPS + DMSO, showing morphological features consistent with reactive astrogliosis, including cell body enlargement, flattening, and increased morphological heterogeneity. Bottom left: Normal + Inflamate® (0.0375 μg/mL), showing morphology comparable to the vehicle control. Bottom right: LPS + Inflamate®, showing partial attenuation of LPS-induced morphological changes, with a greater proportion of cells retaining elongated morphology. Scale bar = 10 μm. These observations are preliminary; further validation using higher-magnification imaging or GFAP immunofluorescence staining is warranted.

## Author Contributions

**Conceptualization**, K.H. and C.-T.T.; **Methodology**, M.K., C.-F.H. and C.-T.T.; **Formal Analysis**, M.K., C.-F.H. and C.-T.T.; **Investigation**, M.K., C.-F.H. and C.-T.T.; **Validation**, M.K., C.-F.H. and C.-T.T.; **Visualization**, M.K., C.-F.H. and C.-T.T.; **Literature Review**, M.K., S.I., C.-F.H., C.-T.T. and T.F.; **Writing—Original Draft Preparation**, M.K. and C.-F.H.; **Writing—Review and Editing**, K.H.; **Project Administration**, K.H., T.F. and S.O.; **Supervision, S.I. and K.H.**; **Funding Acquisition**, M.K. and K.H. All authors have read and agreed to the published version of the manuscript.

*All authors have read and agreed to the published version of the manuscript.

## Data Availability Statement

The data is original and was provided by all authors. Support for the findings of this study is available. Further inquiries can be directed to the corresponding author.

## Acknowledgments

We thank Haruki Uchibori and Yuzuki Takagaki for their assistance in the preparation of various laboratory equipment and materials. We are grateful to RIKEN BioResource Research Center (RIKEN BRC) for providing the mouse AWT astrocyte cell line (RCB5681) through the National Bio-Resource Project of the Ministry of Education, Culture, Sports, Science and Technology (MEXT), Japan. We also extend our gratitude to Kenbics (Japan) for providing Inflamate® to support this research, enabling commercially available dietary supplement products to be evaluated in cellular assays and thereby establishing the developmental potential for future experiments.

## References

1. Ganz, T. and E. Nemeth, Iron homeostasis in host defence and inflammation. Nat Rev Immunol, 2015. 15(8): p. 500–10.

2. Ross, A.C., Impact of chronic and acute inflammation on extra- and intracellular iron homeostasis. Am J Clin Nutr, 2017. 106(Suppl 6): p. 1581S–1587S.

3. Dringen, R., et al., The pivotal role of astrocytes in the metabolism of iron in the brain. Neurochem Res, 2007. 32(11): p. 1884–90.

4. Qian, Z.M. and Y. Ke, Brain iron transport. Biol Rev Camb Philos Soc, 2019. 94(5): p. 1672–1684.

5. Manu, D.R., et al., Astrocyte Involvement in Blood-Brain Barrier Function: A Critical Update Highlighting Novel, Complex, Neurovascular Interactions. Int J Mol Sci, 2023. 24(24).

6. Patani, R., G.E. Hardingham, and S.A. Liddelow, Functional roles of reactive astrocytes in neuroinflammation and neurodegeneration. Nat Rev Neurol, 2023. 19(7): p. 395–409.

7. Rochfort, K.D. and P.M. Cummins, The blood-brain barrier endothelium: a target for pro-inflammatory cytokines. Biochem Soc Trans, 2015. 43(4): p. 702–6.

8. Jager, R., et al., International Society of Sports Nutrition Position Stand: Probiotics. J Int Soc Sports Nutr, 2019. 16(1): p. 62.

9. Li, Y., et al., Ulinastatin suppresses lipopolysaccharide induced neuro-inflammation through the downregulation of nuclear factor-kappaB in SD rat hippocampal astrocyte. Biochem Biophys Res Commun, 2015. 458(4): p. 763–70.

10. Sheng, W., et al., Pro-inflammatory cytokines and lipopolysaccharide induce changes in cell morphology, and upregulation of ERK1/2, iNOS and sPLA(2)-IIA expression in astrocytes and microglia. J Neuroinflammation, 2011. 8: p. 121.

11. Singh, S., et al., Lipopolysaccharide induced altered signaling pathways in various neurological disorders. Naunyn Schmiedebergs Arch Pharmacol, 2022. 395(3): p. 285–294.

12. Yuste, J.E., et al., Implications of glial nitric oxide in neurodegenerative diseases. Front Cell Neurosci, 2015. 9: p. 322.

13. Boullerne, A.I. and J.A. Benjamins, Nitric oxide synthase expression and nitric oxide toxicity in oligodendrocytes. Antioxid Redox Signal, 2006. 8(5-6): p. 967–80.

14. Ageeva, T., A. Rizvanov, and Y. Mukhamedshina, NF-kappaB and JAK/STAT Signaling Pathways as Crucial Regulators of Neuroinflammation and Astrocyte Modulation in Spinal Cord Injury. Cells, 2024. 13(7).

15. Jain, M., et al., Role of JAK/STAT in the Neuroinflammation and its Association with Neurological Disorders. Ann Neurosci, 2021. 28(3-4): p. 191–200.

16. Xin, H., et al., Hydrogen Sulfide Attenuates Inflammatory Hepcidin by Reducing IL-6 Secretion and Promoting SIRT1-Mediated STAT3 Deacetylation. Antioxid Redox Signal, 2016. 24(2): p. 70–83.

17. Fleming, R.E., Iron and inflammation: cross-talk between pathways regulating hepcidin. J Mol Med (Berl), 2008. 86(5): p. 491–4.

18. Verga Falzacappa, M.V., et al., STAT3 mediates hepatic hepcidin expression and its inflammatory stimulation. Blood, 2007. 109(1): p. 353–8.

19. Nemeth, E. and T. Ganz, Hepcidin-Ferroportin Interaction Controls Systemic Iron Homeostasis. Int J Mol Sci, 2021. 22(12).

20. Galy, B., M. Conrad, and M. Muckenthaler, Mechanisms controlling cellular and systemic iron homeostasis. Nat Rev Mol Cell Biol, 2024. 25(2): p. 133–155.

21. Moreira, A.C., G. Mesquita, and M.S. Gomes, Ferritin: An Inflammatory Player Keeping Iron at the Core of Pathogen-Host Interactions. Microorganisms, 2020. 8(4).

22. Vela, D., Hepcidin, an emerging and important player in brain iron homeostasis. J Transl Med, 2018. 16(1): p. 25.

23. Lee, J. and D.H. Hyun, The Interplay between Intracellular Iron Homeostasis and Neuroinflammation in Neurodegenerative Diseases. Antioxidants (Basel), 2023. 12(4).

24. Chen, W., et al., Ferritin in ferroptosis: Implications for neurodegenerative diseases (Review). Int J Mol Med, 2026. 57(5).

25. Santana-Codina, N. and J.D. Mancias, The Role of NCOA4-Mediated Ferritinophagy in Health and Disease. Pharmaceuticals (Basel), 2018. 11(4).

26. Santana-Codina, N., A. Gikandi, and J.D. Mancias, The Role of NCOA4-Mediated Ferritinophagy in Ferroptosis. Adv Exp Med Biol, 2021. 1301: p. 41–57.

27. Guo, W., et al., NCOA4-mediated ferritinophagy promoted inflammatory responses in periodontitis. J Periodontal Res, 2021. 56(3): p. 523–534.

28. Ward, R.J., D.T. Dexter, and R.R. Crichton, Iron, Neuroinflammation and Neurodegeneration. Int J Mol Sci, 2022. 23(13).

29. David, S., et al., Dysregulation of Iron Homeostasis in the Central Nervous System and the Role of Ferroptosis in Neurodegenerative Disorders. Antioxid Redox Signal, 2022. 37(1-3): p. 150–170.

30. Corradini, E., J.L. Babitt, and H.Y. Lin, The RGM/DRAGON family of BMP co-receptors. Cytokine Growth Factor Rev, 2009. 20(5-6): p. 389–98.

31. Fisher, A.L. and J.L. Babitt, Coordination of iron homeostasis by bone morphogenetic proteins: Current understanding and unanswered questions. Dev Dyn, 2022. 251(1): p. 26–46.

32. Key, B. and G.J. Lah, Repulsive guidance molecule A (RGMa): a molecule for all seasons. Cell Adh Migr, 2012. 6(2): p. 85–90.

33. Zhang, J., et al., Repulsive guidance molecules b (RGMb): molecular mechanism, function and role in diseases. Expert Rev Mol Med, 2024. 26: p. e24.

34. Hata, K., et al., RGMa inhibition promotes axonal growth and recovery after spinal cord injury. J Cell Biol, 2006. 173(1): p. 47–58.

35. Hata, K., et al., Signaling mechanisms of axon growth inhibition. Drug News & Perspectives, 2006. 19(9): p. 541–547.

36. Hata, K., et al., Unc5B associates with LARG to mediate the action of repulsive guidance molecule. J Cell Biol, 2009. 184(5): p. 737–50.

37. Liddelow, S.A., et al., Neurotoxic reactive astrocytes are induced by activated microglia. Nature, 2017. 541(7638): p. 481–487.

38. Liddelow, S.A. and B.A. Barres, Reactive Astrocytes: Production, Function, and Therapeutic Potential. Immunity, 2017. 46(6): p. 957–967.

39. Clayton, B.L.L. and S.A. Liddelow, Heterogeneity of Astrocyte Reactivity. Annu Rev Neurosci, 2025. 48(1): p. 231–249.

40. Bretheau, F., et al., The alarmin interleukin-1alpha triggers secondary degeneration through reactive astrocytes and endothelium after spinal cord injury. Nat Commun, 2022. 13(1): p. 5786.

41. Zhang, Q., et al., Blocking C3d(+)/GFAP(+) A1 Astrocyte Conversion with Semaglutide Attenuates Blood-Brain Barrier Disruption in Mice after Ischemic Stroke. Aging Dis, 2022. 13(3): p. 943–959.

42. Wang, S., et al., Transcriptome Analysis Reveals Dynamic Microglial-Induced A1 Astrocyte Reactivity via C3/C3aR/NF-kappaB Signaling After Ischemic Stroke. Mol Neurobiol, 2024. 61(12): p. 10246–10270.

43. Bozic, I., D. Savic, and I. Lavrnja, Astrocyte phenotypes: Emphasis on potential markers in neuroinflammation. Histol Histopathol, 2021. 36(3): p. 267–290.

44. Chang, J., et al., Transplantation of A2 type astrocytes promotes neural repair and remyelination after spinal cord injury. Cell Commun Signal, 2023. 21(1): p. 37.

45. Wang, J., et al., Inhibition of A1 Astrocytes and Activation of A2 Astrocytes for the Treatment of Spinal Cord Injury. Neurochem Res, 2023. 48(3): p. 767–780.

46. Li, H., et al., Targeting astrocytes polarization after spinal cord injury: a promising direction. Front Cell Neurosci, 2024. 18: p. 1478741.

47. Zhang, H., et al., Astrocyte-mediated inflammatory responses in traumatic brain injury: mechanisms and potential interventions. Front Immunol, 2025. 16: p. 1584577.

48. Li, T., et al., Microglia induce the transformation of A1/A2 reactive astrocytes via the CXCR7/PI3K/Akt pathway in chronic post-surgical pain. J Neuroinflammation, 2020. 17(1): p. 211.

49. Thau-Habermann, N., et al., Parthenolide regulates microglial and astrocyte function in primary cultures from ALS mice and has neuroprotective effects on primary motor neurons. PLoS One, 2025. 20(3): p. e0319866.

50. Lawrence, J.M., et al., Roles of neuropathology-associated reactive astrocytes: a systematic review. Acta Neuropathol Commun, 2023. 11(1): p. 42.

51. Eo, S.J. and Y.H. Leem, Effects of exercise intensity on the reactive astrocyte polarization in the medial prefrontal cortex. Phys Act Nutr, 2023. 27(2): p. 19–24.

52. Jiang, T., et al., Physical exercise modulates the astrocytes polarization, promotes myelin debris clearance and remyelination in chronic cerebral hypoperfusion rats. Life Sci, 2021. 278: p. 119526.

53. Peternelj, T.T. and J.S. Coombes, Antioxidant supplementation during exercise training: beneficial or detrimental? Sports Med, 2011. 41(12): p. 1043–69.

54. Li, X., et al., Potential Mechanisms of Exercise-Mediated Ferroptosis Regulation in Central Nervous System Diseases. Mol Neurobiol, 2025. 63(1): p. 198.

55. Peng, C., et al., From bench to bedside, boswellic acids in anti-inflammatory therapy - mechanistic insights, bioavailability challenges, and optimization approaches. Front Pharmacol, 2025. 16: p. 1692443.

56. Taherzadeh, D., et al., Acetyl-11-Keto-β-Boswellic Acid (AKBA) Prevents Lipopolysaccharide-Induced Inflammation and Cytotoxicity on H9C2 Cells. Evid Based Complement Alternat Med, 2022. 2022: p. 2620710.

57. Huang, H., et al., Isolation and characterization of two flavonoids, engeletin and astilbin, from the leaves of Engelhardia roxburghiana and their potential anti-inflammatory properties. J Agric Food Chem, 2011. 59(9): p. 4562–9.

58. Altalbawy, F.M.A., et al., Unlocking potentials of a prenylated flavonoid: Xanthohumol’s role in combating inflammatory conditions and Rhematic diseases. Cytokine, 2026. 198: p. 157092.

59. Azuero, M., C.F. Wenceslau, and W. Tan, Xanthohumol: Mechanistic Actions and Emerging Evidence as a Multi-Target Natural Nutraceutical. Nutrients, 2026. 18(3).

60. Pagliari, S., et al., Antioxidant and Anti-Inflammatory Effect of Cinnamon (Cinnamomum verum J. Presl) Bark Extract after In Vitro Digestion Simulation. Foods, 2023. 12(3).

61. Peng, J., et al., The role and mechanism of cinnamaldehyde in cancer. J Food Drug Anal, 2024. 32(2): p. 140–154.

62. Moustafa, E.M., N.M. Thabet, and K.S. Azab, Boswellic acid disables signal transduction of IL-6-STAT-3 in Ehrlich ascites tumor bearing irradiated mice. Biochem Cell Biol, 2016. 94(4): p. 307–13.

63. Sayed, A.S., et al., Role of 3-Acetyl-11-Keto-Beta-Boswellic Acid in Counteracting LPS-Induced Neuroinflammation via Modulation of miRNA-155. Mol Neurobiol, 2018. 55(7): p. 5798–5808.

64. Sayed, A.S. and N.S. El Sayed, Co-administration of 3-Acetyl-11-Keto-Beta-Boswellic Acid Potentiates the Protective Effect of Celecoxib in Lipopolysaccharide-Induced Cognitive Impairment in Mice: Possible Implication of Anti-inflammatory and Antiglutamatergic Pathways. J Mol Neurosci, 2016. 59(1): p. 58–67.

65. Kostka, M., et al., Muscle-brain crosstalk mediated by exercise-induced myokines - insights from experimental studies. Front Physiol, 2024. 15: p. 1488375.

66. Bustin, S.A., et al., The MIQE guidelines: minimum information for publication of quantitative real-time PCR experiments. Clin Chem, 2009. 55(4): p. 611–22.

67. Urrutia, P.J., D.A. Borquez, and M.T. Nunez, Inflaming the Brain with Iron. Antioxidants (Basel), 2021. 10(1).

68. Fujii, T., et al. RGM Family Involved in the Regulation of Hepcidin Expression in Anemia of Chronic Disease. Immuno, 2024. 4, 266–285 DOI: 10.3390/immuno4030017.

69. Lin, H.Y., The Central Role of BMP Signaling in Regulating Iron Homeostasis, in Bone Morphogenetic Proteins: Systems Biology Regulators, S. Vukicevic and K.T. Sampath, Editors. 2017, Springer International Publishing: Cham. p. 345–356.

70. Zhao, Q., et al., The origins and dynamic changes of C3- and S100A10-positive reactive astrocytes after spinal cord injury. Front Cell Neurosci, 2023. 17: p. 1276506.

71. Hong, Y., et al., High-frequency repetitive transcranial magnetic stimulation improves functional recovery by inhibiting neurotoxic polarization of astrocytes in ischemic rats. J Neuroinflammation, 2020. 17(1): p. 150.

72. Abbott, N.J., L. Ronnback, and E. Hansson, Astrocyte-endothelial interactions at the blood-brain barrier. Nat Rev Neurosci, 2006. 7(1): p. 41–53.

73. El Baassiri, M.G., et al., Pharmacologic Toll-like receptor 4 inhibition skews toward a favorable A1/A2 astrocytic ratio improving neurocognitive outcomes following traumatic brain injury. J Trauma Acute Care Surg, 2023. 95(3): p. 361–367.

74. Li, B., et al., The role of reactive astrocytes in neurotoxicity induced by ultrafine particulate matter. Sci Total Environ, 2023. 867: p. 161416.

75. Giovannoni, F. and F.J. Quintana, The Role of Astrocytes in CNS Inflammation. Trends Immunol, 2020. 41(9): p. 805–819.

76. Che, D.N., et al., Luteolin and Apigenin Attenuate LPS-Induced Astrocyte Activation and Cytokine Production by Targeting MAPK, STAT3, and NF-kappaB Signaling Pathways. Inflammation, 2020. 43(5): p. 1716–1728.

77. Yang, X., et al., The role of the JAK2-STAT3 pathway in pro-inflammatory responses of EMF-stimulated N9 microglial cells. J Neuroinflammation, 2010. 7: p. 54.

78. Yin, J., et al., The role of TLR4/NF-kB signaling axis in pneumonia: from molecular mechanisms to regulation by phytochemicals. Naunyn Schmiedebergs Arch Pharmacol, 2025. 398(11): p. 14559–14588.

79. Liu, C., et al., 6-Bromoindirubin-3’-Oxime Suppresses LPS-Induced Inflammation via Inhibition of the TLR4/NF-kappaB and TLR4/MAPK Signaling Pathways. Inflammation, 2019. 42(6): p. 2192–2204.

80. Lai, J.L., et al., Indirubin Inhibits LPS-Induced Inflammation via TLR4 Abrogation Mediated by the NF-kB and MAPK Signaling Pathways. Inflammation, 2017. 40(1): p. 1–12.

81. Xie, C., et al., Anti-inflammatory Activity of Magnesium Isoglycyrrhizinate Through Inhibition of Phospholipase A2/Arachidonic Acid Pathway. Inflammation, 2015. 38(4): p. 1639–48.

82. Calixto, J.B., M.F. Otuki, and A.R. Santos, Anti-inflammatory compounds of plant origin. Part I. Action on arachidonic acid pathway, nitric oxide and nuclear factor kappa B (NF-kappaB). Planta Med, 2003. 69(11): p. 973–83.

83. Vivancos, M. and J.J. Moreno, Role of Ca(2+)-independent phospholipase A(2) and cyclooxygenase/lipoxygenase pathways in the nitric oxide production by murine macrophages stimulated by lipopolysaccharides. Nitric Oxide, 2002. 6(3): p. 255–62.

84. Choi, S.H., S. Aid, and F. Bosetti, The distinct roles of cyclooxygenase-1 and -2 in neuroinflammation: implications for translational research. Trends Pharmacol Sci, 2009. 30(4): p. 174–81.

85. Abe, A., et al., The role of lysosomal phospholipase A2 in the catabolism of bis(monoacylglycerol)phosphate and association with phospholipidosis. J Lipid Res, 2024. 65(7): p. 100574.

86. Yan, M., et al., SOCS modulates JAK-STAT pathway as a novel target to mediate the occurrence of neuroinflammation: Molecular details and treatment options. Brain Res Bull, 2024. 213: p. 110988.

87. Baker, B.J., L.N. Akhtar, and E.N. Benveniste, SOCS1 and SOCS3 in the control of CNS immunity. Trends Immunol, 2009. 30(8): p. 392–400.

88. Huang, S.N., et al., Aspirin increases ferroportin 1 expression by inhibiting hepcidin via the JAK/STAT3 pathway in interleukin 6-treated PC-12 cells. Neurosci Lett, 2018. 662: p. 1–5.

89. Angmo, S., et al., Identification of Guanosine 5’-diphosphate as Potential Iron Mobilizer: Preventing the Hepcidin-Ferroportin Interaction and Modulating the Interleukin-6/Stat-3 Pathway. Sci Rep, 2017. 7: p. 40097.

90. Abdukarimov, N., K. Kokabi, and J. Kunz, Ferroptosis and Iron Homeostasis: Molecular Mechanisms and Neurodegenerative Disease Implications. Antioxidants (Basel), 2025. 14(5).

91. Levi, S., et al., Iron imbalance in neurodegeneration. Mol Psychiatry, 2024. 29(4): p. 1139–1152.

92. Xu, Y., et al., Astrocyte hepcidin ameliorates neuronal loss through attenuating brain iron deposition and oxidative stress in APP/PS1 mice. Free Radic Biol Med, 2020. 158: p. 84–95.

93. Zhang, X., et al., Hepcidin overexpression in astrocytes alters brain iron metabolism and protects against amyloid-beta induced brain damage in mice. Cell Death Discov, 2020. 6(1): p. 113.

94. Dmytriv, T.R., K.B. Storey, and V.I. Lushchak, Intestinal barrier permeability: the influence of gut microbiota, nutrition, and exercise. Front Physiol, 2024. 15: p. 1380713.

95. Henningsen, K., I. Martinez, and R.J.S. Costa, Exertional Stress-induced Pathogenic Luminal Content Translocation - Friend or Foe? Int J Sports Med, 2024. 45(8): p. 559–571.

96. Pervaiz, N. and L. Hoffman-Goetz, Immune cell inflammatory cytokine responses differ between central and systemic compartments in response to acute exercise in mice. Exerc Immunol Rev, 2012. 18: p. 142–57.

97. Nash, D., et al., IL-6 signaling in acute exercise and chronic training: Potential consequences for health and athletic performance. Scand J Med Sci Sports, 2023. 33(1): p. 4–19.

98. Vargas, N.T. and F. Marino, A neuroinflammatory model for acute fatigue during exercise. Sports Med, 2014. 44(11): p. 1479–87.

99. Svenningsson, P., et al., p11 and its role in depression and therapeutic responses to antidepressants. Nat Rev Neurosci, 2013. 14(10): p. 673–80.

100. Seo, J.S. and P. Svenningsson, Modulation of Ion Channels and Receptors by p11 (S100A10). Trends Pharmacol Sci, 2020. 41(7): p. 487–497.

101. Sofroniew, M.V., Molecular dissection of reactive astrogliosis and glial scar formation. Trends Neurosci, 2009. 32(12): p. 638–47.

102. Gaojian, T., et al., Parthenolide promotes the repair of spinal cord injury by modulating M1/M2 polarization via the NF-kappaB and STAT 1/3 signaling pathway. Cell Death Discov, 2020. 6(1): p. 97.

103. Burda, J.E. and M.V. Sofroniew, Reactive gliosis and the multicellular response to CNS damage and disease. Neuron, 2014. 81(2): p. 229–48.

104. Rasmussen, P., et al., In humans IL-6 is released from the brain during and after exercise and paralleled by enhanced IL-6 mRNA expression in the hippocampus of mice. Acta Physiol (Oxf), 2011. 201(4): p. 475–82.

105. de Souza, R.F., et al., Ultra-Endurance Associated With Moderate Exercise in Rats Induces Cerebellar Oxidative Stress and Impairs Reactive GFAP Isoform Profile. Front Mol Neurosci, 2020. 13: p. 157.

106. Escartin, C., et al., Reactive astrocyte nomenclature, definitions, and future directions. Nat Neurosci, 2021. 24(3): p. 312–325.

107. Moulson, A.J., et al., Diversity of Reactive Astrogliosis in CNS Pathology: Heterogeneity or Plasticity? Front Cell Neurosci, 2021. 15: p. 703810.

